# Temporal multi-modal single-cell analyses reveal dynamic interactions of CAR-T cells with glioblastoma and targeting of antigen-negative neoplastic cells

**DOI:** 10.1101/2024.10.03.616537

**Authors:** Daniel Y. Zhang, Xin Wang, Yusha Sun, Qi Cui, Ernest Nelson, Zhijian Zhang, Brian Huang, Josephine Giles, Radhika Thokala, Daniel R. Connolly, Fadi Jacob, E John Wherry, Timothy Lucas, H. Isaac Chen, Yanhong Shi, Steven Brem, Zev A. Binder, MacLean P. Nasrallah, Ryan D. Salinas, Donald M. O’Rourke, Guo-li Ming, Hongjun Song

## Abstract

CAR-T therapy is a promising new immunotherapy for cancers, but its efficacy for solid tumors requires improvement. A detailed understanding of the interplay between solid tumors and CAR-T cells is critical. Here we report temporal, multi-modal, single-cell profiling of patient-derived glioblastoma organoids with CAR-T treatment. We found that all tumor cell types responded to CAR-T cell activation and contributed to an initially anti-tumor, but subsequently pro-tumor and immune-inhibitory microenvironment, accompanied by CAR-T cell exhaustion. Unexpectedly, CAR-T treatment attenuated glioma stem-like states of both antigen-positive and antigen-negative neoplastic cells and reduced their proliferation via diffusible factors, including IFNγ. Analysis of samples from additional patients, including those in clinical trials, supported these findings. Our study reveals the dynamic interplay among different tumor cells and T cells in adaptive responses to immunotherapy and identifies previously unappreciated benefits of CAR-T therapy directly on antigen-negative neoplastic cells that may be leveraged to enhance therapeutic efficacy.

## INTRODUCTION

Glioblastoma (GBM) remains an incurable and rapidly lethal cancer, and the limited improvements in long-term patient survival over the past two decades call for new treatment strategies. Advances in immune therapies have shown great promise for some cancers, such as remarkable results from CD19 targeted chimeric antigen receptor T (CAR-T) cell therapy for B-cell malignancies^1–4^. Recently, three clinical trials of CAR-T cell therapies for GBM, including those targeting IL13RA2^5^, EGFRvIII^6^, or both IL13RA2/EGFR^7^, have reported exciting initial anti-tumor effects, despite eventual disease recurrence in many cases. These studies, while promising, suggest that improving clinical efficacy in GBM requires novel and fine-tuned approaches that better address the unique biology of this cancer.

Many challenges facing CAR-T cell therapy for GBM have been identified, including heterogeneous expression of the target antigen and an evolving complex immunosuppressive tumor microenvironment (TME)^8–11^. The biological processes underlying these challenges are rooted in an adaptive tumor response from both neoplastic and non-neoplastic cells within the TME; thus, a better understanding of the dynamic interactions between tumor and CAR-T cells is needed to improve therapeutic efficacy. Importantly, as it remains difficult to identify antigen targets that are expressed both ubiquitously and selectively in tumor cells across a broad patient population, and as CAR-T cell therapy often ends up with relapses arising from antigen^-^ neoplastic cells^12^, methods to broaden the anti-tumor effect of CAR-T cell therapy^7^, especially against antigen^-^ neoplastic cells, are critical.

The discovery of cellular plasticity in GBM tumor cells raises the possibility of altering cell states as a new therapeutic strategy^13,14^. Although different models have been proposed for the cellular composition and lineage hierarchy in GBM, there is agreement that neoplastic cells reside along continuums of cell states and exhibit dynamic plasticity throughout tumorigenesis and in response to perturbation^15–17^. Glioma stem cells, although yet to be fully defined^18^, demonstrate enhanced plasticity and have the capacity to generate progeny in different cell states^14,19^. Overall, these processes result in the observed tumor heterogeneity and treatment resistance, but also present opportunities to drive neoplastic cells towards therapeutically advantageous cell states with reduced proliferation and weakened adaptive potential^13^. Furthermore, the TME and individual local niches provide critical signals for cell state identity and transitions^20,21^, and these components, in addition to cell intrinsic pathways, are potential targets for intervention.

One major bottleneck to gain a better understanding of dynamic interactions between GBM and CAR-T cells is the limited availability of patient samples for temporal analysis. Patient-derived GBM organoids (GBOs) have been established as an experimental model that captures both inter- and intra-tumoral heterogeneity and maintains native cell-cell interactions and the TME^22–25^. GBOs contain a diversity of neoplastic cells with varied antigen expression and somatic mutations, and resident non-neoplastic cells, including immune cells, such as macrophages and microglia. In a recent phase 1 clinical trial of dual-targeting CAR-T cells against EGFR and IL13Rα2 in patients with recurrent glioblastoma^7^ (NCT05168423), we performed parallel analyses of six sets of paired patient-derived GBOs treated with the same patient CAR-T cell products. Our data demonstrated an excellent correlation between several measurements of responses in GBOs *in vitro* and in paired patients, including the degree of cytolysis of tumor cells in GBOs with the amount of CAR-T cell engraftment detected in patients’ cerebrospinal fluid (CSF) and cytokine release patterns in GBOs and in patient CSF samples over time (see accompanying manuscript). These results suggest that patient-derived GBOs could serve as high fidelity avatars to address key questions related to how cellular heterogeneity contributes to CAR-T therapy responses, such as how different cell types remodel the immune-related TME, whether antigen^+^ and antigen^-^ cells respond similarly or differently to treatment, and how neoplastic cell states and glioma stem cell niches may be affected by immune activity.

Here we investigated temporal dynamics of CAR-T cell therapy with GBOs using a multi-modal single-cell assay that simultaneously analyzed gene expression, cell surface protein expression, and somatic genetic mutations within the same cell. We found large changes in the transcriptomic states that were temporally linked and reciprocally driven by CAR-T cell activation. The tumor adaptive responses dramatically altered the cell-cell interaction network with both anti- and pro-tumor properties, involving not only neoplastic cells and CAR-T cells, but also immune and brain parenchymal components of the TME. Unexpectedly, we discovered changes in neoplastic cell niche signaling that attenuated glioma stem-like states and reduced the proliferation of both antigen^+^ and antigen^-^ neoplastic cells. We validated some of these key findings using pre- and short-term post-CAR-T cell treatment patient tumor samples from a clinical trial (NCT02209376)^26^. In addition, our mechanistic studies identified that these effects were mediated by diffusible factors, including IFNɣ. Our study provides a rich resource revealing the dynamic evolution of the tumor and TME during CAR-T cell treatment and identifies novel means to harness immune activity for GBM treatment.

## RESULTS

### Temporal Multi-modal Single-cell Analyses of CAR-T Cell Treatment in a GBO Model

We used three different CAR constructs, including CAR-T19 (“CD19”), which is used clinically to target B cells^1^ as a negative control, “2173”, which targets a tumor-specific EGFRvIII mutant and has been used in clinical trials for GBM^26,27^, and “806”, which was derived from a monoclonal antibody with high affinity to the EGFRvIII mutant and lower affinity to wild-type and other variants of EGFR^28,29^ and was one of dual targeting CARs in the recent clinical trial^7^. For single-cell analyses, we used GBOs from five patients with different initial EGFRvIII expression levels, including one patient without any detectable expression as a control (Figure S1B-C and Table S1). To avoid potential differences in T cell properties from different patients, we engineered T cells from the same donor for different CARs and applied them to all GBOs from different patients for comparison. The co-culture system demonstrated three features. First, it yielded robust T cell engraftment and expansion as well as prominent tumor cell death during the initial period, especially in GBOs with high EGFRvIII levels (Figure 1A-B, S1B-C and Supplementary Movie 1). Second, it exhibited graded immune activity, as measured by IFNɣ production, depending on the initial levels of EGFRvIII in the GBOs (Figure S1B-D). Third, by day 14, EGFRvIII^+^ neoplastic cells and CD3^+^ T cells co-existed with reduced cell death and diminished IFNɣ production in GBOs (8167), indicating incomplete antigen loss, tumor persistence, and decreased T cell activity (Figure 1B-C and S1D). Notably, primary GBM samples from the same patient (8167) who participated in a clinical trial of 2173 CAR-T cell therapy (NCT02209376)^26^ similarly demonstrated incomplete antigen loss in the presence of persistent CD3^+^ T cell infiltration after treatment (Figure 1C). Together, these results indicate that our co-culture system can model the initial tumor responses and subsequent resistance of GBM during CAR-T cell treatment.

**Figure 1.**
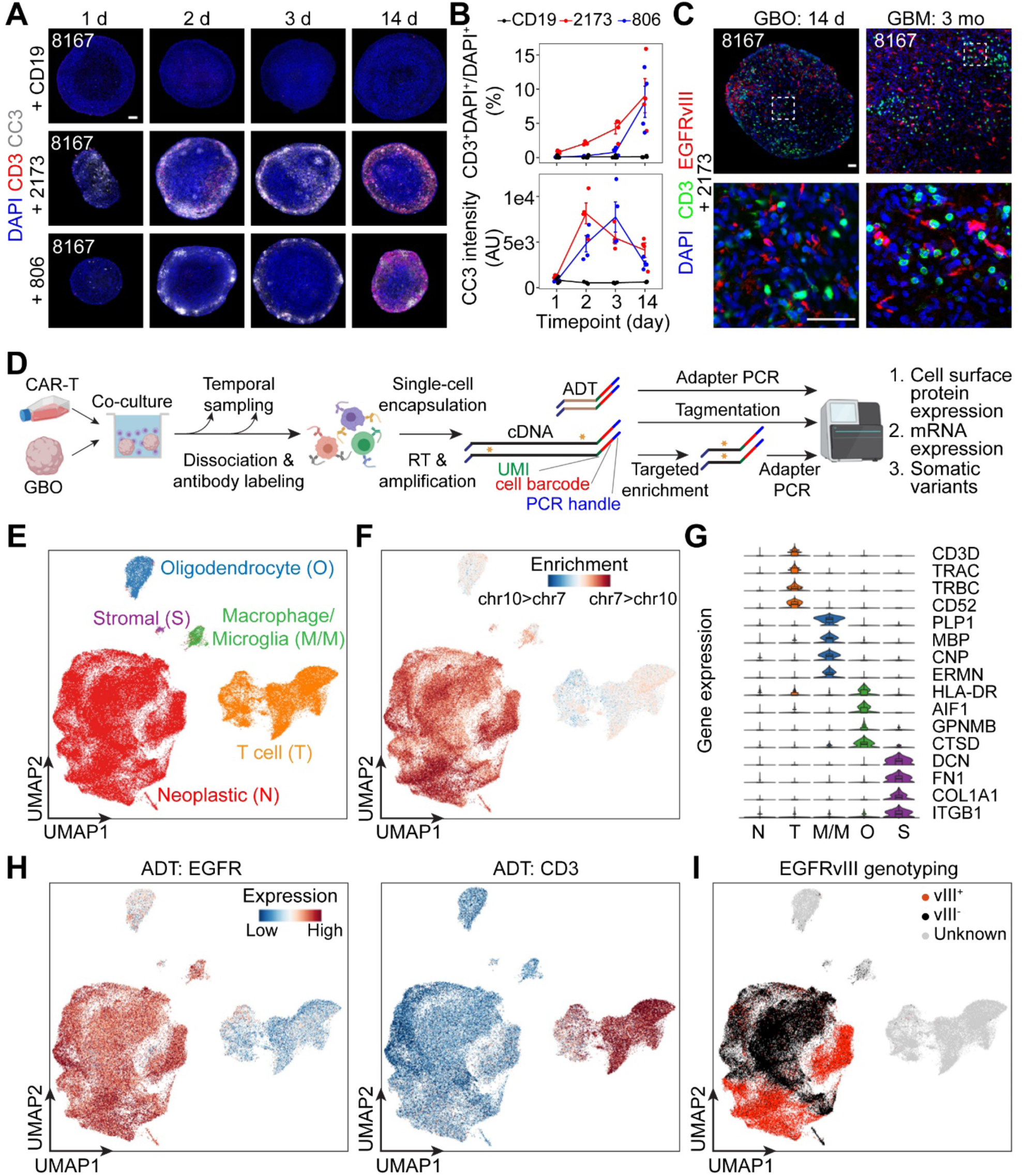
Multi-modal, temporal, single-cell analyses in a GBO and CAR-T cell co-culture model. (A-B) Sample confocal images of CD3 and cleaved caspase 3 (CC3) immunostaining of UP-8167 GBOs co-cultured with CD19, 2173, or 806 CAR-T cells at 1, 2, 3, and 14 days (A; Scale bar: 100 μm) and quantification of the percentage of CD3^+^ cells among DAPI^+^ cells in GBOs and of the CC3 immunostaining intensity (B). Individual data points are plotted with error bars corresponding to mean ± SEM (n = 4). (C) Sample confocal images of CD3 and EGFRvIII immunostaining of UP-8167 GBOs co-cultured with 2173 CAR-T cells for 14 days and of the same patient’s tumor 3 months after receiving 2173 CAR-T cell treatment with concomitant Pembrolizumab, a PD-1 inhibitor. Scale bars: 50 μm. (D) Schematic diagram depicting the temporal sampling of the GBO and CAR-T cell co-cultures followed by multi-modal single-cell analyses for the whole transcriptome expression by scRNA-seq, cell surface protein expression of selected targets by CITE-seq, and somatic variant genotyping, with all information for the same individual cells. (E) UMAP plot of the aggregated scRNA-seq data for all GBOs from 5 patients and at all 5 time points (0, 1, 2, 3, and 14 days). Major cell types are highlighted. (F) UMAP plot of the aggregated scRNA-seq data colored by enrichment for genes located on chromosome 7 vs. chromosome 10. Note that amplification of chromosome 7 and deletion of chromosome 10 indicates neoplastic cells. (G) Violin plot of the aggregated scRNA-seq data showing marker gene enrichment corresponding to the assigned cell types. (H) UMAP plot of the aggregated scRNA-seq data colored by cell surface protein expression of EGFR and CD3 analyzed using antibody-derived tags (ADTs) by CITE-seq. (I) UMAP plot of the aggregated scRNA-seq data colored by EGFRvIII genotyping assignment. Also see **Figure S1**, **Table S1**, and **Supplementary Movie 1**.

In addition to single-cell RNA-sequencing (scRNA-seq) on the Drop-seq platform^30^, we performed simultaneous CITE-seq^31^ using antibody-derived tags for 15 cell surface proteins (Figure 1D and Table S2) to examine protein expression levels, which in some cases yielded higher sensitivity than scRNA-seq. In the scRNA-seq dataset (Figure 1E and S1E), neoplastic cells were identified by the presence of copy number aberrations (CNAs; Figure S1F), especially Chr7 amplification and Chr10 deletion^32^ (Figure 1F). Different types of non-neoplastic cells were broadly identified by the absence of CNAs and enrichment of known markers, including oligodendrocytes (O), macrophages/microglia (M/M), stromal cells (S), and T cells (Figure 1E and 1G). The surface protein expression levels were consistent with cell type assignments as EGFR was highly expressed in neoplastic cells and CD3 was expressed only in T cells (Figure 1H).

From the same full-length cDNA library, we used the Genotyping of Transcriptomes (GoT) method^33^ to identify EGFRvIII mutant transcripts (Figure 1D and Figure S1G). There were several challenges with this particular target, including the distant location of the mutation from the cell barcode, the large size of the exon 2-7 deletion in EGFRvIII, and the heterogeneity of the extended 3’ UTR in the captured transcripts (Figure S1G-H). Optimization of circularization GoT included adaption of an oligo-dA primer to capture the 3’ end of all transcripts regardless of 3’ UTR structures and a combination of primers complementary to both exon 2 and 8 to obtain final sequencing libraries of similar sizes from both EGFRvIII^-^ and EGFRvIII^+^ transcripts, respectively (Figure S1I-K). Ultimately, we achieved a high genotyping efficiency, and the amplification and overexpression of EGFR in neoplastic cells allowed us to set stringent criteria for confident genotyping, which was concordant with our characterization of these samples by immunohistology (Figure 1I and S1C, S1L-O).

Together, our multi-modal approach provides a comprehensive characterization of individual cells for combined transcriptomic profiling, identification of neoplastic and non-neoplastic cells, surface protein expression levels of selected targets, and EGFRvIII mutation status.

### Concomitant Shifts in Cell States in All Cell Types upon CAR-T Cell Treatment

We performed temporal multi-modal single-cell analyses of GBOs from five patients in response to three different CAR-T cell treatments with sampling on days 0, 1, 2, 3 and 14 (Table S2). In controls with CD19 or EGFRvIII^-^ GBOs (8539), T cells were very low in abundance after medium change on day 1 with essentially no engraftment and expansion, and correspondingly, the gene expression of tumor cell populations remained largely unchanged throughout the treatment (Figure 2A and S2A-D).

**Figure 2.**
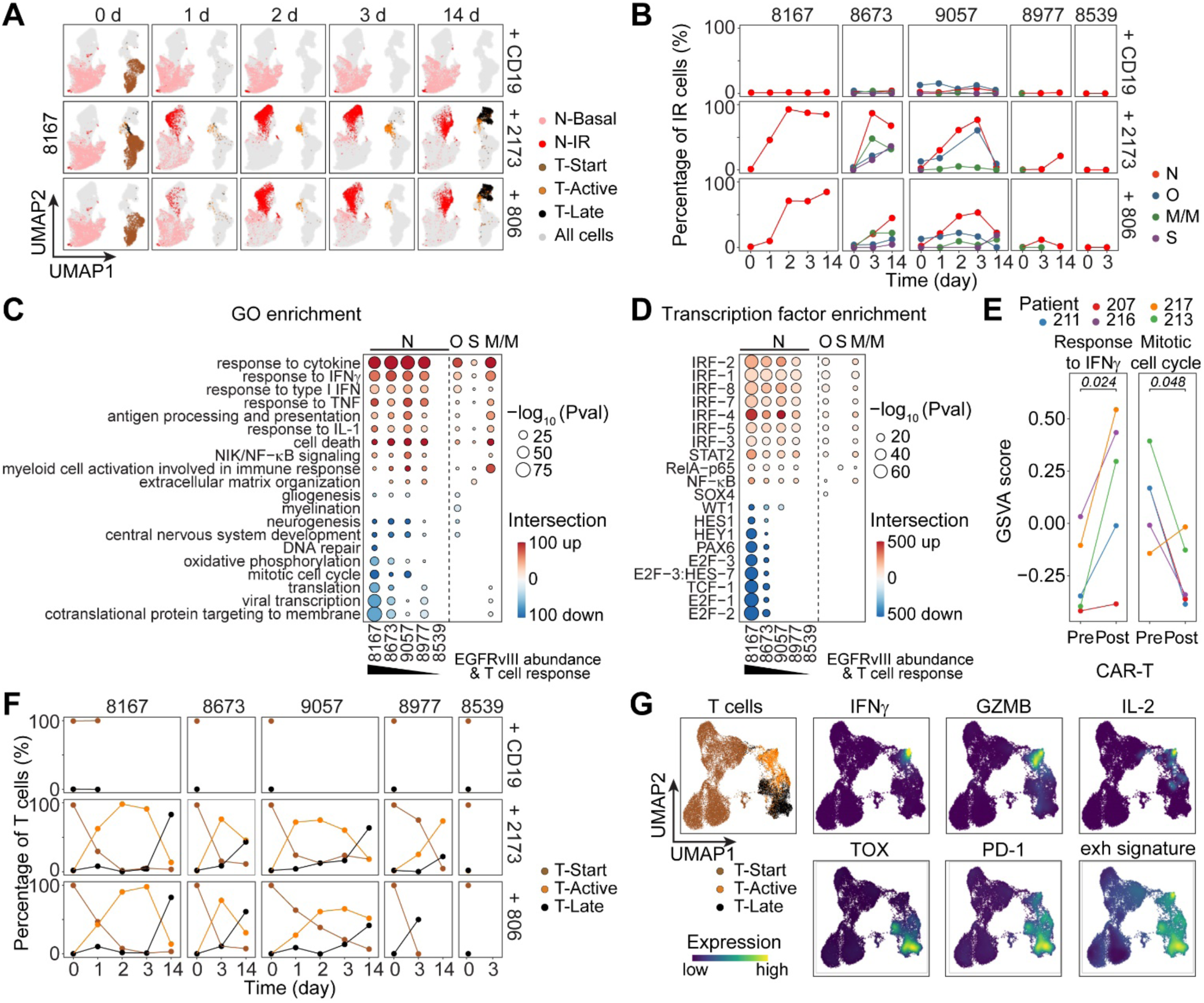
Time-dependent shift in cell states in co-cultured GBOs and CAR-T cells. (A) UMAP plots of scRNA-seq data of UP-8167 GBOs co-cultured with CD19, 2173, and 806 CAR-T cells at different time points. Basal and immune responsive (IR) states for neoplastic cells (N) and start, active, and late states for T cells are indicated by different colors. See similar UMAP plots for GBOs from 4 other patients in Figures S2A-D and aggregated data in Figure S2E. (B) Quantification of the percentage of different tumor cell types in IR states for neoplastic cells (N), oligodendrocytes (O), macrophages/microglia (M/M), and stromal cells (S). The plots are ordered from left to right with decreasing initial EGFRvIII levels in GBOs from 5 different patients. (C-D) Bubble plots of biological process GO term (C) and transcription factor (D) enrichment of up- and down-regulated genes in the IR state for neoplastic cells from GBOs of different patients and for combined non-neoplastic cells from GBOs of all patients. The size of the bubble corresponds to the - log_10_P-value and the color depicts the number of genes intersecting with the pathway. (E) GSVA scores of response to IFNɣ and mitotic cell cycle signatures in bulk RNA-seq data from tumor samples of patients pre- and post-treatment with 2137 CAR-T cells in a clinical trial. The two signatures are from^95^. P-values were calculated using a paired student’s t-test. (F) Quantification of the percentage of T cells in different states in co-culture with GBOs from 5 patients at different time points in response to 3 different CAR-T cells. (G) UMAP density plots of sub-clustered aggregated T cells in different states after SAVER-X denoising. Shown are plots for expression of markers for T cell activation (IFNɣ, GZMB and IL-2) and T cell exhaustion (TOX, PD-1), as well as the broad T cell exhaustion signature from^38^. Also see **Figure S2** and **Tables S2, S3** and **S4**.

For EGFRvIII^+^ GBOs co-cultured with 2173 and 806, there were rapid shifts in gene expression of all tumor cells, including both neoplastic and non-neoplastic cells, from the basal state towards what we broadly termed “immune responsive” (IR) states, as identified by unsupervised clustering analyses (Figure 2A and S2A-E). The extent and durability of these changes correlated with the initial EGFRvIII levels in the GBOs (Figure 2B and S1C). Compared to basal states, IR states shared similar transcriptomic signatures among GBOs from different patients and among different tumor cell types (Table S3), including upregulation of cytokine response pathways to IFNɣ, type I interferon, TNF⍺, and IL-1 (Figure 2C), and increased expression of target genes of interferon response factor (IRF), STAT, and NF-κB family transcription factors (Figure 2D). These IRF regulons, especially those of IRF1 and IRF8, are important in myeloid development and have been found to contribute to a stable transcriptional program hijacked by GBM to help evade immune attack^34,35^. Upregulation of the response to IFNɣ persisted through 14 days even though IFNɣ levels had decreased to baseline (Figure S1D and S2F). As expected for a diffusible factor, both EGFRvIII^+^ and EGFRvIII^-^ neoplastic cells exhibited strong responses to IFNɣ (Figure S2G). In contrast to the shared upregulated genes, downregulated genes, pathways, and transcription factor activities varied across different GBOs and tumor cell types. Neoplastic cells showed downregulation of genes related to neural development, cell cycle, and metabolic pathways, as well as target genes of the cell cycle-promoting E2F family, Notch pathway downstream factors HES1 and HEY1, and neural stem cell-related PAX6. Downregulation of these genes was specific to neoplastic cells and the magnitude of downregulation was correlated with initial EGFRvIII levels in GBOs (Figure 2C-D).

To corroborate our results in the GBO and CAR-T cell co-culture model, we performed bulk RNA-seq of paired tumor samples before and shortly after treatment from five patients with GBM who participated in a previous 2173 CAR-T cell therapy clinical trial (NCT02209376)^26^ (Table S1). In the post-treatment samples, there was also upregulation of cytokine response pathways and increased expression of T cell marker genes CD3D, CD4, and CD8A, indicating elevated T cell abundance, as well as downregulation of the mitotic cell cycle signature (Figure 2E; S2H-I and Table S4).

Concordant with dynamic gene expression changes in tumor cells, T cells from co-cultures of EGFRvIII^+^ GBOs with 2173 and 806 also exhibited a shift in the cell state as identified by unsupervised clustering analysis (Figure 2A, 2F and S2A-E). The T cell active state was observed at early time points with the expression of classical markers, such as IFNɣ, GZMB, and IL-2 (Figure 2G), and was temporally correlated with a shift into the IR states of tumor cells (Figure 2B and 2F). The T cell late state exhibited upregulation of TOX, a transcriptional regulator associated with T cell exhaustion^36,37^, PD-1, an immune checkpoint inhibitor, and general T cell exhaustion-associated gene expression signatures^38^ (Figure 2G and Table S4).

Together, these results show a cascade of cell state changes initiated by CAR-T cell activation upon encountering its target antigen, leading to T cell cytokine secretion, reciprocal cytokine responses across all tumor cell types, and T cell exhaustion with tumor persistence.

### Unbiased Cell-Cell Interaction Analysis between T Cells and Tumor Cells of Different States

Building on the observation that dynamic cell state changes in co-cultured CAR-T cells and GBOs are driven by reciprocal interactions, we performed a broad and unbiased cell-cell interaction inference analysis to gain a more comprehensive insight into these processes. We used CellPhoneDB, which includes a curated ligand-receptor database and an established method to identify statistically significant interactions^39,40^. We first focused on the interactions between T cells and tumor cells in different states (Figure 3A and Table S5). Across different neoplastic and non-neoplastic cell states, the number of cell-cell interactions was higher in the active T cell state than in the start or late T cell states (Figure 3B). In addition, the number of interactions with T cells was greater in the IR state than in the basal state for neoplastic cells, but not for non-neoplastic cell types (Figure 3B). Following previous approaches to use gene expression of the ligand(s) and receptor(s) as a measurement of interaction strength^39,40^, we also found that most changes in interactions between cells in different states were due to the emergence or loss of interactions, rather than strengthening or weakening of pre-existing interactions (Figure 3C and S3A). The exception was with macrophages/microglia, where several pre-existing interactions with T cells in the basal state were strengthened in the IR state, including T cell suppressive interactions between OPN (SPP1) and CD44^41^ (Figure 3C and S3B-C).

**Figure 3.**
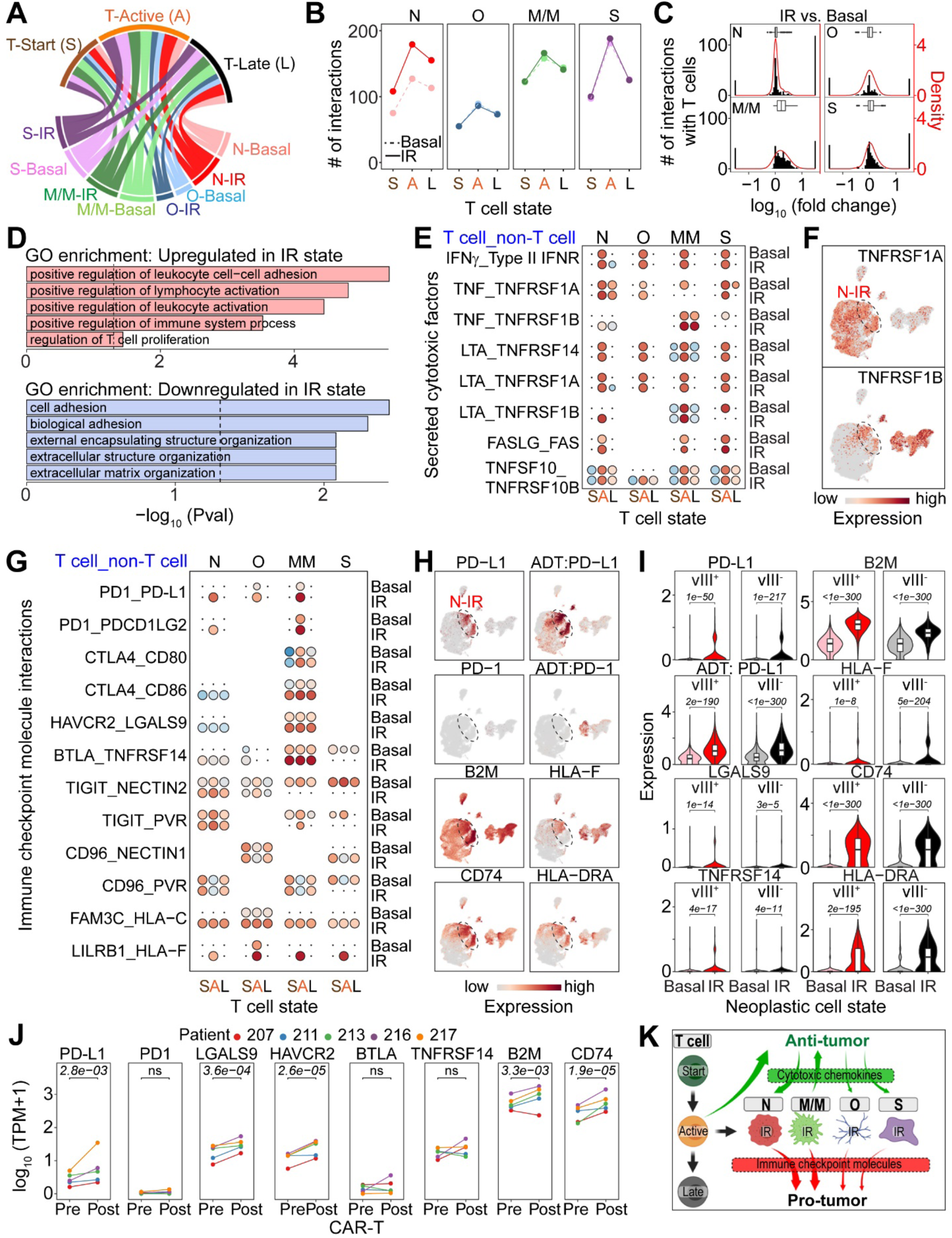
Cell-cell interaction inference analysis between T cells and tumor cells in different states. (A) Chord diagram depicting inferred cell-cell interactions between different tumor cell populations at basal or immune responsive (IR) states and T cells at different states (S: Start; A: Active; L: Late). Width of chords is scaled with the number of statistically significant cell-cell interactions. N: neoplastic cells; O: oligodendrocyte; M/M: macrophages/microglia; S: stromal cells. (B) Quantification of the number of cell-cell interactions identified in different tumor cell states and different T cell states. (C) Histogram of the log_10_fold-change of the interaction strength between IR vs. basal states of different tumor cells with T cells. Interactions that are statistically significant only in basal or IR states are plotted arbitrarily at -1.5 and 1.5, respectively, for visualization. Red line shows a kernel density fit. Inset shows a boxplot summarizing the distribution of these differential interactions. Note that the histogram and box plots center around 0 for N, O and S, but shift to the right for M/M, indicating that many T cell interactions with M/M were strengthened in their IR state, whereas they were either newly present or lost in the IR states of N, O, or S cells. (D) Barplots showing GO enrichment terms for upregulated or downregulated interactions between T cells and IR state tumor cells. Dashed vertical line indicates the significance cutoff of P-value < 0.05. (E) Interaction bubble plots of secreted cytotoxic factors. Note increased interaction strength with T cells in their active state for different tumor cell types at both their basal and IR states. For a detailed description of these interaction bubble plots, see Figure S3E. (F) UMAP plot showing expression of TNFR1 (TNFRSF1A) and TNFR2 (TNFRSF1B) in the aggregated scRNA-seq dataset. The neoplastic cells in the IR state are highlighted by the dashed line circle. Note that TNFR1 is expressed broadly in neoplastic cells, whereas TNFR2 expression is largely limited to the IR state. (G) Interaction bubble plot of immune inhibitory molecules from different tumor cell types and corresponding receptors in T cells. Note that the interaction strength was increased with tumor cells in their IR states. (H) UMAP plot showing expression of example immune inhibitory molecules at the mRNA and cell surface protein (ADT) levels as indicated. (I) Violin plot of expression of immune inhibitory molecules in IR and basal state neoplastic cells separated by the EGFRvIII genotype. Only neoplastic cells with 2173 CAR-T cell treatment were included in the analysis. P values are calculated using the Wilcoxon rank sum test with Bonferonni correction for multiple hypothesis testing. (J) Gene expression of immune inhibitory molecules in pre- and post-CAR-T cell treatment patient samples from the 2173 trial based on bulk RNA-seq. Significance testing was performed using DESeq2 with Benjamini-Hochberg correction for multiple testing. (K) A summary drawing depicting changes in T cell state where active T cells induce “immune responsive” (IR) states across all tumor cells. These IR tumor cells, largely neoplastic (N) cells and macrophage/microglia (M/M) are engaged in both anti-tumor and pro-tumor interactions through cytotoxic cytokines and inhibitory immune checkpoint molecules, respectively. The width of arrow indicates relative contribution. Also see **Figure S3** and **Tables S1** and **S5**.

We developed a matrix-style interaction bubble-plot to visualize the strength of specific interactions among various cell types in different states using the IFNɣ and IFNɣ receptor (type II IFNR) interaction as an example (Figure S3D-E). The IFNɣ and IFNɣ receptor interactions were present between active T cells and all neoplastic and non-neoplastic tumor cells in both their basal and IR states. The specificity of this interaction was driven by the selective expression of IFNɣ in the active T cell state, whereas the IFNɣ receptor was expressed consistently throughout at both mRNA and protein levels (Figure S3F).

We identified biological processes that were differentially enriched in interactions between T cells and tumor cells in the basal versus IR states (Figure 3D). Notably, cell adhesion pathways were enriched in both up- and down-regulated interactions, suggesting remodeling of T cell adhesion interactions in response to immune activity (Figure 3D). For example, interactions of tumor cell receptor ICAM1 with T cell integrins, including LFA-1 (integrin aLb2 complex, consisting of a heterodimer of ITGAL and ITGB2), were upregulated in the IR state of neoplastic cells and macrophages/microglia (Figure S3I-J). Similar observations were made in post-treatment patient samples from the 2173 trial, where ICAM1, ITGAL, and ITGAX were upregulated (Figure S3K). In contrast, T cell L-selectin (SELL) and semaphorin 4D (SEMA4D) interactions were downregulated across most tumor cell types (Figure S3I-J). Both these up- and downregulated pathways are associated with T cell trafficking and migration^42–44^, and these changes may promote T cell engraftment within GBOs.

We also observed interactions between secreted TNF family cytotoxic factors by active T cells, including TNF⍺, Lymphotoxin Alpha (LTA), Fas Ligand (FASLG), and TRAIL (TNFSF10), and corresponding death receptors in tumor cells (Figure 3E and S3G). We validated the expression of TNFα in T cells by immunohistology (Figure S3H). In addition to active T cells, IR-state macrophages/microglia also upregulated TNF⍺ levels (Figure S3G). Receptor expression was largely unchanged between basal and IR states with the notable exception of TNFR2 (TNFRSF1B). Neoplastic cells expressed only TNFR1 (TNFRSF1A) in the basal state, but expressed TNFR2 in the IR state (Figure 3F). Post-treatment patient samples from the 2173 trial also showed upregulation of TNF⍺ and TNFR2, whereas TNFR1 expression levels were unchanged (Figure S3K). TNFR1 and TNFR2 signaling are known to exhibit varying effects on neural stem cells and immune modulatory activity, and induced TNFR2 signaling in neoplastic cells has been suggested to play an important role in the adaptive tumor response^45,46^.

This cell-cell interaction analysis allowed us to identify the sources of various upregulated immune inhibitory molecules in the IR states of different tumor cell types^47–49^ (Figure 3G). PD-L1 (CD274) was highly upregulated in the IR states of both neoplastic cells and macrophages/microglia, and correspondingly, PD-1 (PDCD1) was elevated in the active and late states of T cells (Figure 3G-H). TIM3 (HAVCR2) ligand LGALS9 and BTLA ligand HVEM (TNFRSF14) were expressed higher in IR state macrophages/microglia, whereas CTLA4 ligands (CD80 and CD86) were expressed only in IR state macrophages/microglia (Figure 3G). TIGIT ligand CD112 (NECTIN2) was prominently upregulated in IR state neoplastic cells (Figure 3G). Furthermore, neoplastic cells in the IR state upregulated MHC I and II subunit genes B2M and CD74 (Figure 3G-H). Some non-classical HLA molecules, such as HLA-F and HLA-G, were expressed in IR state tumor cells, which can engage receptors, such as LILRB1, to co-opt immune tolerance pathways and inhibit immune activity^50,51^ (Figure 3G-H and Table S3). Upregulation of MHC II in cancers has been proposed to promote antigen presentation to CD4^+^ T cells and contribute to antitumor immunity; however, they may also interact with LAG3 to suppress overall T cell activity^52,53^. All of these molecules were expressed by both EGFRvIII^+^ and EGFRvIII^-^ neoplastic cells (Figure 3I), indicating that antigen^-^ cells contribute similarly to the adaptive immune response and suggesting that upregulation of these immune inhibitory molecules is likely linked to the production of diffusible factors. Post-treatment patient samples from the 2173 trial also showed a similar upregulation of many immune checkpoint inhibitors, including PD-L1, LGALS9, and HAVCR2 (Figure 3J).

Together, these results indicate that CAR-T cell treatment leads to initial anti-tumor responses, including signals from multiple tumor cell types to promote T cell engraftment and upregulate secreted TNF family cytotoxic molecules, followed by pro-tumor responses through the upregulation of an array of immune inhibitory molecules, mostly from neoplastic cells and macrophages/microglia (Figure 3K).

### CAR-T Cell Treatment Alters Ligand-receptor Expression that Can Regulate Recruitment and Differentiation of Peripherally-derived Leukocytes

Next, we investigated how CAR-T cell treatment may impact ligand-receptor signaling that regulates the recruitment of immune cells to tumors. Both EGFRvIII^+^ and EGFRvIII^-^ IR state neoplastic cells were the primary source of CXCR3 ligands (CXCR9/10/11), whereas the CCR4 ligand CCL5 and the CXCR4 ligand CXCL12 were also expressed by IR state macrophages/microglia and stromal cells (Figure 4A-B and Figure S4A). These signals function both individually and cooperatively to enhance the recruitment of CAR-T and host T cells into solid tumors^54–57^. IR state macrophages/microglia also upregulated CCR5 ligands (CCL3 and CCL4) (Figure S4B), which play an important role in the recruitment of T cells^58,59^. Similarly, many of these chemokines and/or their receptors were upregulated in post-treatment patient samples from the 2173 trial, including CXCL9, CXCR3, CCL5, CCR4, and CXCL12 (Figure 4C). These results suggest that the induced expression of these chemoattractants by neoplastic and multiple non-neoplastic cells in the TME upon CAR-T cell activation may further recruit additional T cells to exit the circulation and enter the tumor parenchyma.

**Figure 4.**
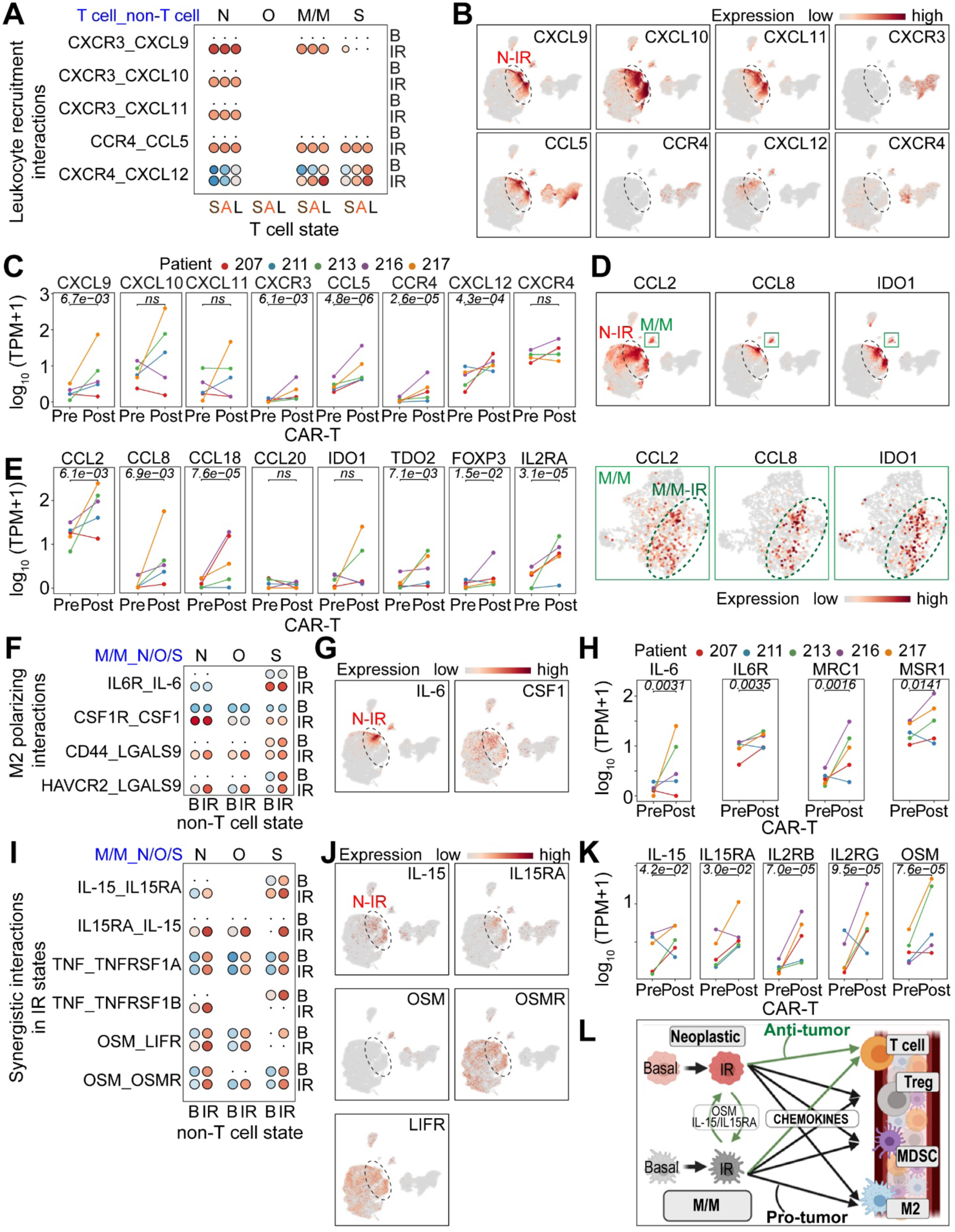
CAR-T cell treatment induces changes of ligand-receptor expression that can impact recruitment and differentiation of peripheral leukocytes. (A-C) Upregulation of T cell recruitment interactions in IR states of different tumor cell types. Shown in (A) is an interaction bubble plot. Shown in (B) are UMAP plots of scRNA-seq data colored by expression of both chemokines and corresponding receptors. Neoplastic cells in the IR state are highlighted with the dashed line circle. Shown in (C) are plots of gene expression of chemokine and receptor molecules in pre- and post-CAR-T cell treatment patient samples from the 2173 trial. Significance testing was performed using DESeq2 and Benjamini-Hochberg correction for multiple hypothesis testing. (D-E) Upregulation of Treg recruitment chemokines and differentiation promoting metabolic enzymes in the IR states of different tumor cells (D) and in samples from the 2173 trial along with additional Treg markers, FOXP3 and CD25 (IL2RA) (E). Similar as in (B-C). Also shown in (D) are UMAP plots of subclustered macrophages/microglia with the IR state highlighted with a dashed line circle (See Figure S3G). (F-H) Upregulation of M2 polarizing cytokines in IR states of different tumor cells, as shown in an interaction bubble plot (F), UMAP plots of scRNA-seq data colored by gene expression (G), and samples from the 2173 trial along with additional markers of M2 polarization, CD206 (MRC1) and CD204 (MSR1) (H). Similar as in (A-C). (I-K) Additively stronger anti-tumor interactions between neoplastic cells and macrophage/microglia when both are in IR states, as shown in an interaction bubble plot (I), UMAP plots of scRNA-seq data colored by gene expression (J), and samples from the 2173 trial (K). Similar as in (A-C). (L) A summary drawing depicting both pro-tumor and anti-tumor immune cell recruitment through secreted chemokines from neoplastic cells (N) and macrophages/microglia (M/M). Additionally, anti-tumor interactions between N and M/M are highlighted. Also see **Figures S3** and **S4**.

In parallel with anti-tumor leukocyte recruitment signals, pro-tumor signals that recruit and differentiate leukocytes with immunosuppressive functions, such as Regulatory T cells (Tregs), M2-like macrophages, and Myeloid-derived Suppressor Cells (MDSCs), were also upregulated in IR states. Chemokines CCL2, CCL8, CCL18, and CCL20, were produced by IR state EGFRvIII^+^ and EGFRvIII^-^ neoplastic cells and macrophages/microglia (Figure 4D and S4C-D). We also validated CCL2 expression in the treated tumor cells by immunohistology (Figure S3H). These factors have been shown to recruit Tregs in solid tumors, leading to suppressed immune activity and further tumor growth^60–65^. Within the TME, CCL20 produced by IR state neoplastic cells can stimulate macrophages/microglia to produce CCL2^60^. IR state neoplastic cells and macrophages/microglia also upregulated metabolic enzymes IDO1 and TDO (Figure 4D and S4C-D), which can inhibit T cell activity and promote Treg differentiation through local depletion of tryptophan and production of kynurenine^66–68^. Post-treatment patient samples from the 2173 trial also showed upregulated Treg-recruiting chemokines CCL2, CCL8 and CCL18 and Treg marker genes FOXP3 and CD25 (IL2RA) (Figure 4E).

IR state tumor cells also generated signals that function to accumulate immunosuppressive M2-like macrophages and MDSCs. Some chemokines associated with Treg recruitment, such as CCL2, CCL8, and CCL18, can also recruit inhibitory myeloid cells^60,69,70^. In addition, IR state EGFRvIII^+^ and EGFRvIII^-^ neoplastic cells, macrophages/microglia, and stromal cells all produced IL-6 (Figure 4F-G and S4E), which promotes M2 polarization and MDSC recruitment and amplification^71^. The M2 polarizing factors CSF1 and LGALS9^72–74^ were also upregulated in the IR state, largely in neoplastic cells (Figure 4F-G and S4E). Post-treatment patient samples from the 2173 trial showed similar upregulation of IL-6 and two markers of M2 polarization, MRC1 (CD206) and MSR1 (CD204) ^75^ (Figure 4H).

Macrophages/microglia are highly abundant resident immune cells in GBM, and there is growing appreciation for their interactions with neoplastic cells in tumorigenesis and treatment resistance^76^. Interestingly, several interactions became stronger in the IR states of both cell types, suggesting opportunities for enhancing the existing anti-tumor factors. IL-15 and IL15RA were both upregulated in the IR states of macrophages/microglia and interacting neoplastic cells (Figure 4I-J). The IL-15/IL15RA complex acts as a potent pro-inflammatory stimulant to NK and T cells, and augmentation of this response may help further increase immune cell activity in the tumor^77^. Macrophage/microglia-derived Oncostatin M (OSM) and the corresponding receptors OSMR and LIFR in neoplastic cells were also cumulatively stronger when both cell types were in the IR states (Figure 4I-J). OSM signaling in GBM has been correlated with increased cytotoxic T cell activity and may enhance tumor susceptibility to immune attack^78^. Our results indicate that OSM signaling itself may be enhanced by CAR-T cell treatment, suggesting a positive feedback loop driven by this therapeutic approach. Post-treatment patient samples from the 2173 trial also showed upregulation of IL-15, IL15RA, and OSM (Figure 4K).

Taken together, these results highlight the complex evolution of cell-cell interactions upon CAR-T cell treatment, some of which support antitumor activity, whereas others promote pro-tumor resistance through immune inhibition (Figure 4L). The TME plays a critical role in the adaptive tumor response, with substantial contributions from both neoplastic and non-neoplastic cells, resulting in an overall phenotype of tumor persistence and eventual CAR-T cell exhaustion. In particular, macrophages/microglia play a unique role, both independently and cooperatively with neoplastic cells, and some of these interactions suggest opportunities for augmenting immune activity in the tumor.

### CAR-T Cell Treatment Downregulates Glioma Stem Cell Signatures and Reduces Proliferation across Neoplastic Cells

Given that our GBO model maintains native cell-cell interactions as in patient tumors^22^, we also examined changes in interactions among neoplastic cells upon CAR-T cell treatment. We subclustered neoplastic cells and performed ligand-receptor inference analysis between different subpopulations. We identified several interactions that were consistently enriched or weakened in the IR state across EGFRvIII^+^ GBOs from different patients (Figure 5A and S5A). Some of the enriched interactions involve MHC I and II molecules, NRG1, and DPP4, which proteolytically processes many secreted molecules, including CXCL9/10/11, and CCL5, to regulate and tune their activities^79^. A similar upregulation was observed in post-treatment patient samples from the 2173 trial (Figure S5B).

**Figure 5.**
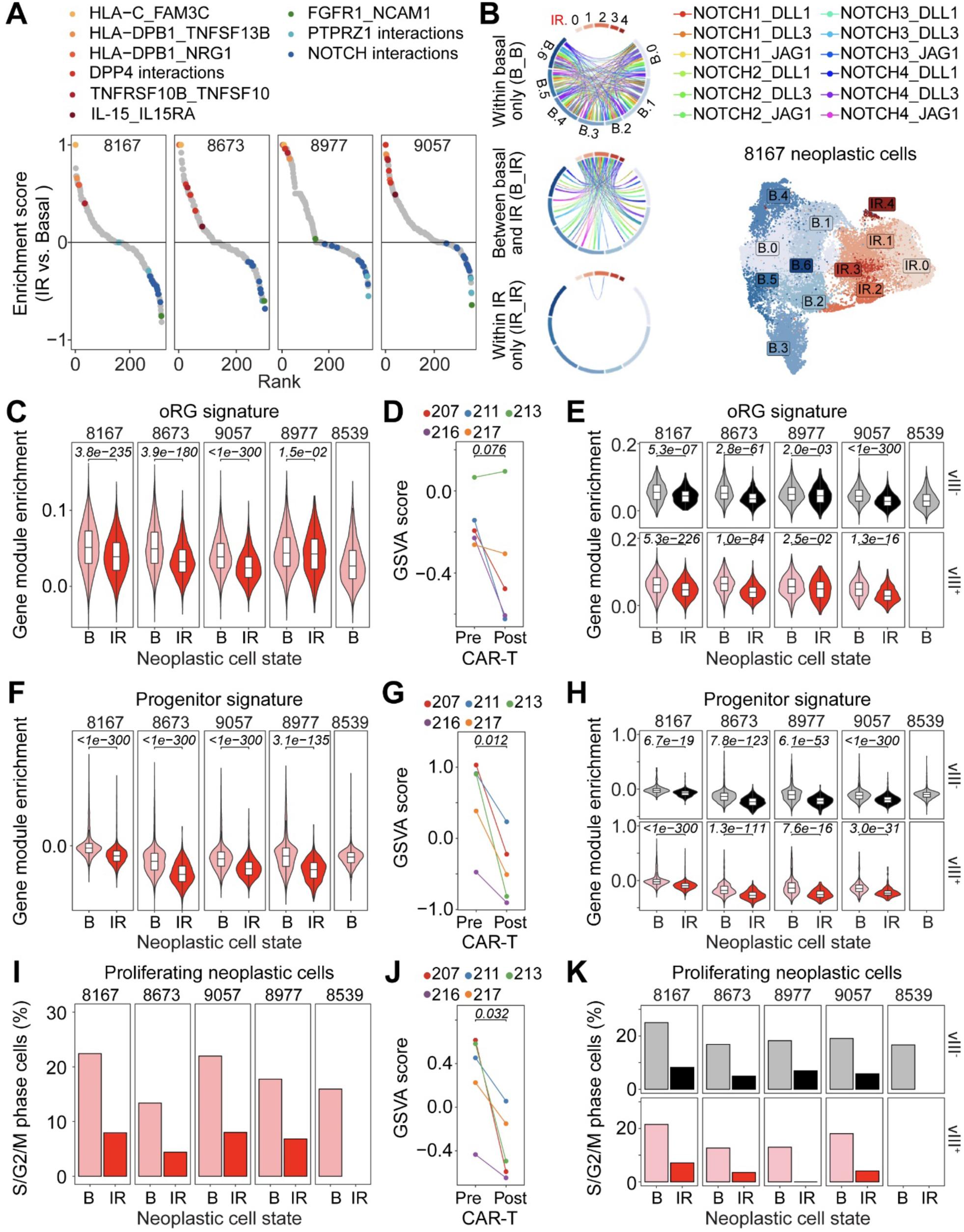
CAR-T cell treatment induces changes of cell-cell interactions within the neoplastic cell population. (A) Cell-cell interactions within the neoplastic cell populations in EGFRvIII^+^ GBOs from 4 patients rank ordered by enrichment in IR vs. basal states. Selected shared IR state-enriched (red dots) and basal state-enriched (blue dots) interactions and interaction families are highlighted. (B) Chord diagram of Notch interactions within basal state populations, between basal and IR populations, and within IR populations. Also shown is the UMAP plot of sub-clustered neoplastic cells in UP-8167, including 6 clusters for the basal state (B.0-B.5) and 5 clusters for the IR state (R.0-R.4). (C, F) Enrichment of oRG (C) and progenitor (F) signatures in the basal vs. IR state of neoplastic cells. The oRG signature is from^83^ and the progenitor signature is from^84^. P-values from a Wilcoxon rank sum test are shown. (D, G) GSVA score of oRG (D) and progenitor (G) signatures in pre- and post-CAR-T cell treatment patient samples from the 2173 trial. P-values were calculated using a paired student’s t-test. (E, H) Enrichment for oRG (E) and progenitor (H) signatures in the IR vs. basal state of neoplastic cells separated by the EGFRvIII genotype. Only cells from the 2173 CAR-T cell treatment were included in this analysis. Similar as in (C and F). (I) Quantification of the percentage of S, G2, and M-phase cells in the basal vs. IR state among neoplastic cells. (J) GSVA score of cell cycle signatures in pre- and post-CAR-T cell treatment patient samples from the 2173 trial. The cell cycle signature is the same as used to determine S/G2/M phase cells in (I). Similar as in (E). (K) Quantification of the percentage of S, G2, and M-phase cells in the basal vs. IR state among neoplastic cells separated by the EGFRvIII genotype. Also see **Figure S5** and **Table S4**.

Interestingly, notable stem cell-associated interactions were downregulated, including those involving FGFR1, PTPRZ1, and Notch^80–82^ (Figure 5A). A recent study suggested that PTPRZ1 is a critical mediator of the outer radial glia (oRG)-like glioma stem cell state, which contributes to the cellular heterogeneity and locally invasive behavior of GBM ^83^. Both the ligand (PTN) and receptor (PTPRZ1) were downregulated in IR state neoplastic cells, leading to downstream signaling changes mimicking those found in previously reported PTPRZ1 knockdowns in both patient-derived GBM cells and fetal human brain cells^83^ (Figure S5C-D). The Notch family is composed of four receptors, which were all expressed in GBOs, and five ligands, of which DLL1, DLL3, and JAG1 were expressed (Figure S5E). There were complex interactions involving Notch ligands and receptors in the basal state, which were decreased between basal and IR state cells, and were nearly abolished for cells in the IR state (Figure 5B and S5G). Both Notch ligands and receptors were downregulated in the IR state, but the loss of Notch ligands, which were expressed in more distinct populations, appeared to be a greater driving force than the loss of Notch receptors, which were expressed uniformly (Figure S5E-F). There was also downregulation of the Notch signaling pathway in IR state neoplastic cells in GBOs and in post-treatment patient samples from the 2173 trial (Figure S5H-I).

The finding of reduced PTPRZ1 and Notch signaling led us to examine changes in the glioma stem-like properties of neoplastic cells in basal versus IR states. We found that the oRG-like glioma stem cell signature^83^ was consistently decreased in the IR state of both EGFRvIII^+^ and EGFRvIII^-^ neoplastic cells across GBOs from different patients (Figure 5C and 5E). Both EGFRvIII^+^ and EGFRvIII^-^ IR state neoplastic cells also exhibited downregulation of a glioma progenitor signature identified by an independent method^84^ (Figure 5F, 5H and S5J, S5L). In post-treatment samples from the 2173 trial, there was a decrease in the glioma progenitor signature for all five samples and a decrease in the oRG signature for four out of the five samples (Figure 5D, 5G and S5K). Importantly, there was also decreased proliferation of both EGFRvIII^+^ and EGFRvIII^-^ neoplastic cells in the IR state in GBOs across patients as well as cell cycle signatures in post-treatment patient samples from the 2173 trial (Figure 5I-K).

Together, these results show that CAR-T cell treatment leads to downregulation of glioma stem cell signatures in both EGFRvIII^+^ and EGFRvIII^-^ neoplastic cells, accompanied by an overall reduction in cell proliferation. These analyses reveal an unexpected direct therapeutic effect on neoplastic cells lacking the CAR-T target antigen, thus potentially broadening the potential efficacy of cellular immunotherapy.

### Diffusible Factors from CAR-T Cell Treatment Downregulate Glioma Stem Cell Signatures and Reduce Proliferation of Neoplastic Cells

The similar impact of CAR-T cell treatment on both EGFRvIII^+^ and EGFRvIII^-^ neoplastic cells suggests the involvement of diffusible factors in downregulating glioma stem cell signatures and reducing cell proliferation. To directly examine the role of diffusible factors and confirm our findings with a large cohort of independent patient samples, we collected conditioned media from EGFRvIII^+^ GBOs co-cultured with CD19 (CD19M) or 806 CAR-T cells (806M) to treat GBOs derived from 27 patient samples with diverse somatic mutations (Table S1). We performed bulk RNA-seq 3 days after treatment and PCA showed a shift in the same rightward direction along PC1 from CD19M to 806M for GBOs from all 27 patients (Figure 6A). Differential gene expression analysis identified extensive changes, including upregulation of PD-L1 and CXCR3 ligands as well as downregulation of glioma stem cell markers PTPRZ1 and PROM1 (CD133)^18,85^ and proliferation marker Ki67 (Figure 6B and Table S6). Compared to the up- and downregulated genes in IR state neoplastic cells from our scRNA-seq, we found that exposure to 806M alone recapitulated many of the gene expression and biological pathway changes observed in the co-culture of GBOs and CAR-T cells (Figure 6C-D and S6A-B). However, 806M treatment alone could not fully recapitulate all features of the IR state, such as the downregulation of some metabolic processes (Figure 6D).

**Figure 6.**
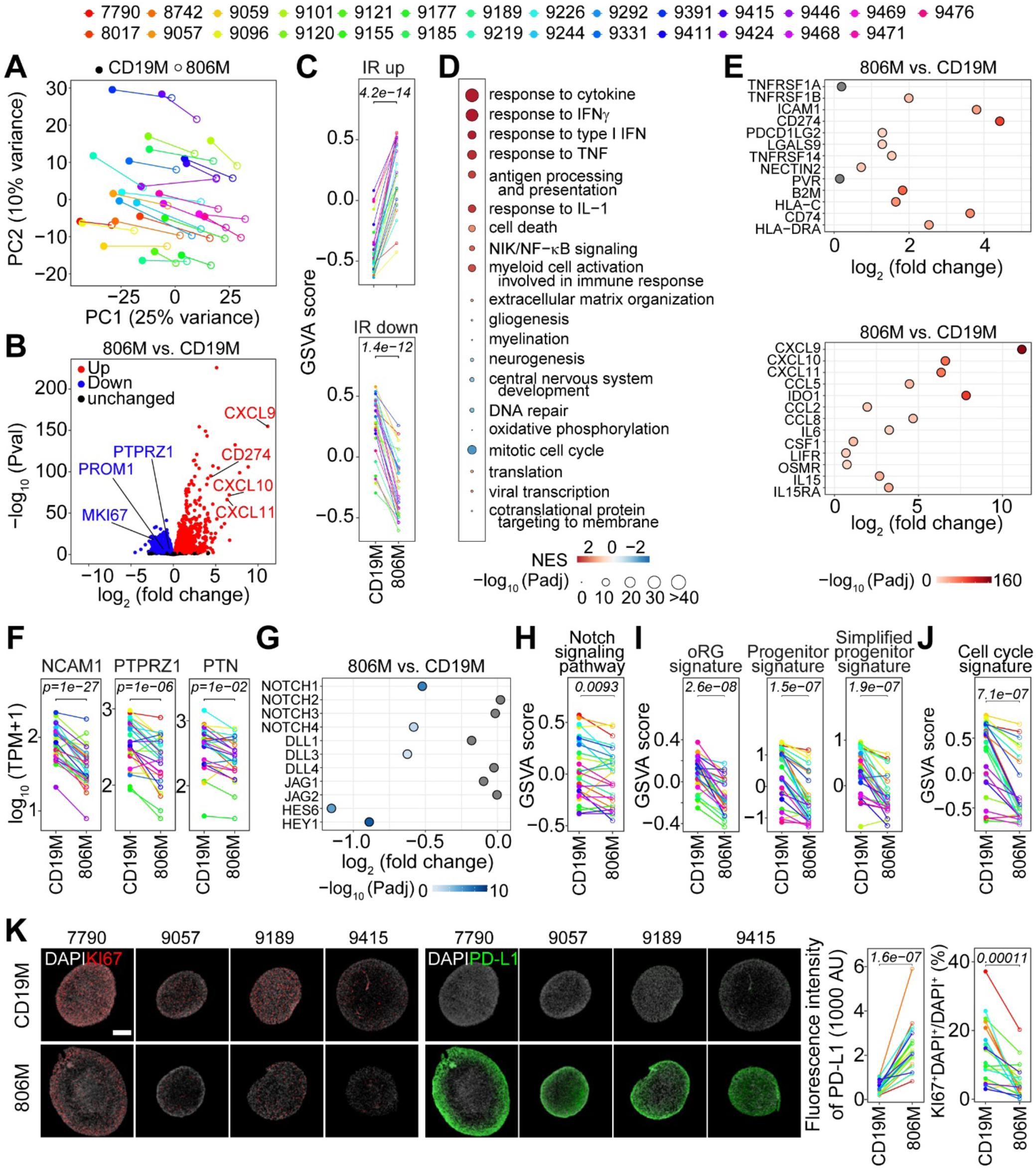
Many of the neoplastic cell phenotypes in the GBO and CAR-T cell co-culture are mediated by diffusible factors. (A) PCA plot of bulk RNA-seq data of GBOs derived from 27 patient samples treated with CD19 (CD19M) or 806 (806M) conditioned media for 3 days. Note the right shift of 806M along PC1 for all samples. (B) Volcano plot of gene expression for 806M vs. CD19M. Cut off for significance was set at an adjusted P-value < 0.01. Note that chemokine genes (CXCL9, CXCL10 and CXCL11) and immune inhibitory molecule CD274 (PD-L1) were upregulated, whereas glioma stem cell-associated genes PTPRZ1 and PROM1 (CD133) and proliferation marker (MKI67) were downregulated. (C) GSVA scores of IR signatures in CD19M or 806M treated GBOs derived from 27 patient samples. The IR up and IR down signatures were from scRNA-seq of GBOs in co-culture with CAR-T cells as in Figure 2C. P-values were calculated using a paired student’s t-test. (D) Bubble plot showing GSEA results of up- and down-regulated pathways found in the neoplastic IR state (same terms as in Figure 2C) for 806M vs. CD19M treatments. Point size corresponds to - log_10_P-value and color corresponds to the normalized enrichment score (NES). (E) Scatterplot of upregulated ligand and receptor molecules associated with immune inhibitory interactions (top panel) and leukocyte recruitment and differentiation signals (bottom panel). X-axis indicates the log_2_fold-change of gene expression for 806M vs. CD19M, and point color indicates the - log_10_P-value. (F) Expression of downregulated neoplastic niche interaction genes, NCAM1, PTPRZ1, and PTN in GBOs derived from 27 patient samples treated with either CD19M or 806M. P-values were calculated using a paired student’s t-test. (G) Scatterplot of expression of ligands, receptors, and downstream molecules of the Notch signaling pathway. Similar as in (E) (H-J) GSVA scores of Notch signaling pathway (H), glioma stem cell (I), and cell cycle (J) signatures in GBOs derived from 27 patient samples treated with CD19M and 806M. P-values were calculated using a paired student’s t-test. (K) Sample confocal images of KI67 and PD-L1 immunostaining of GBOs treated with CD19M or 806M. Scale bar: 200 μm. (J) Quantification of PD-L1 immunostaining intensity and percentage of KI67^+^ cells in CD19M and 806M treated GBOs derived from 20 patient samples. P values are calculated with a paired student’s t-test. See Figure S6D for plots of data for GBOs from individual patient samples. Also see **Figure S6** and **Tables S1** and **S6**.

Exposure to 806M also recapitulated the upregulation of many ligands and receptors that we identified in scRNA-seq to be altered in the cell-cell interaction networks with T cells and macrophages/microglia, including the TNFR2 and OSM receptors, LIFR and OSMR (Figure 6E and S6C). Many changes in ligands and receptors involved in cell-cell interactions among neoplastic cells were also recapitulated with 806M, including downregulation of NCAM1, PTPRZ1, PTN, some members of the Notch family ligands, Notch targets HEY1 and HES6, and the Notch pathway in general (Figure 6F-H). Importantly, glioma stem cell and cell cycle signatures were downregulated with 806M alone (Figure 6I-J). Immunohistology confirmed the upregulation of PD-L1 expression and decreased percentages of KI67^+^ cells in GBOs with 806M (Figure 6K and S6D) as well as with conditioned medium from EGFRvIII^+^ GBOs co-cultured with 2173 (2173M) for a subset of GBOs tested (Figure S6E).

To examine the impact of diffusible factors directly on glioma stem cells, we exposed four established patient-derived glioma stem cell lines^86^ to CD19M or 806M and found drastically reduced cell growth and formation of tumor spheres from single cells with 806M compared with CD19M, indicating reduced glioma stem cell growth and stemness (Figure S6F).

Together, these results demonstrate that diffusible factors can recapitulate many features of direct CAR-T cell treatment, particularly the downregulation of glioma stem cell signatures and reduction of tumor cell proliferation.

### IFNɣ is Required, but not Sufficient, for Conditioned Medium-mediated Downregulation of Glioma Stem Cell Signatures and Reduced Tumor Cell Proliferation

To identify factors in the conditioned media upon CAR-T cell treatment that are important in mediating downregulation of glioma stem cell signatures and reduction of tumor cell proliferation, we performed a cytokine array analysis of 806M in comparison to CD19M. Many of the upregulated secreted factors identified by scRNA-seq, including CXCR3 ligands (CCL5, CXCL12), IL-6 and CSF1, were more abundant in 806M, confirming our scRNA-seq findings at the protein level (Figure 7A and Table S7). IFNɣ was one of the most upregulated proteins (Figure 7A and S7A) and IFNɣ signaling was also one of the top upregulated cytokine response pathways enriched in IR states (Figure 2C). In addition, IFNɣ signaling has been shown to regulate neural stem cells^87^ and to reduce glioma stem cell growth ^88^.

**Figure 7.**
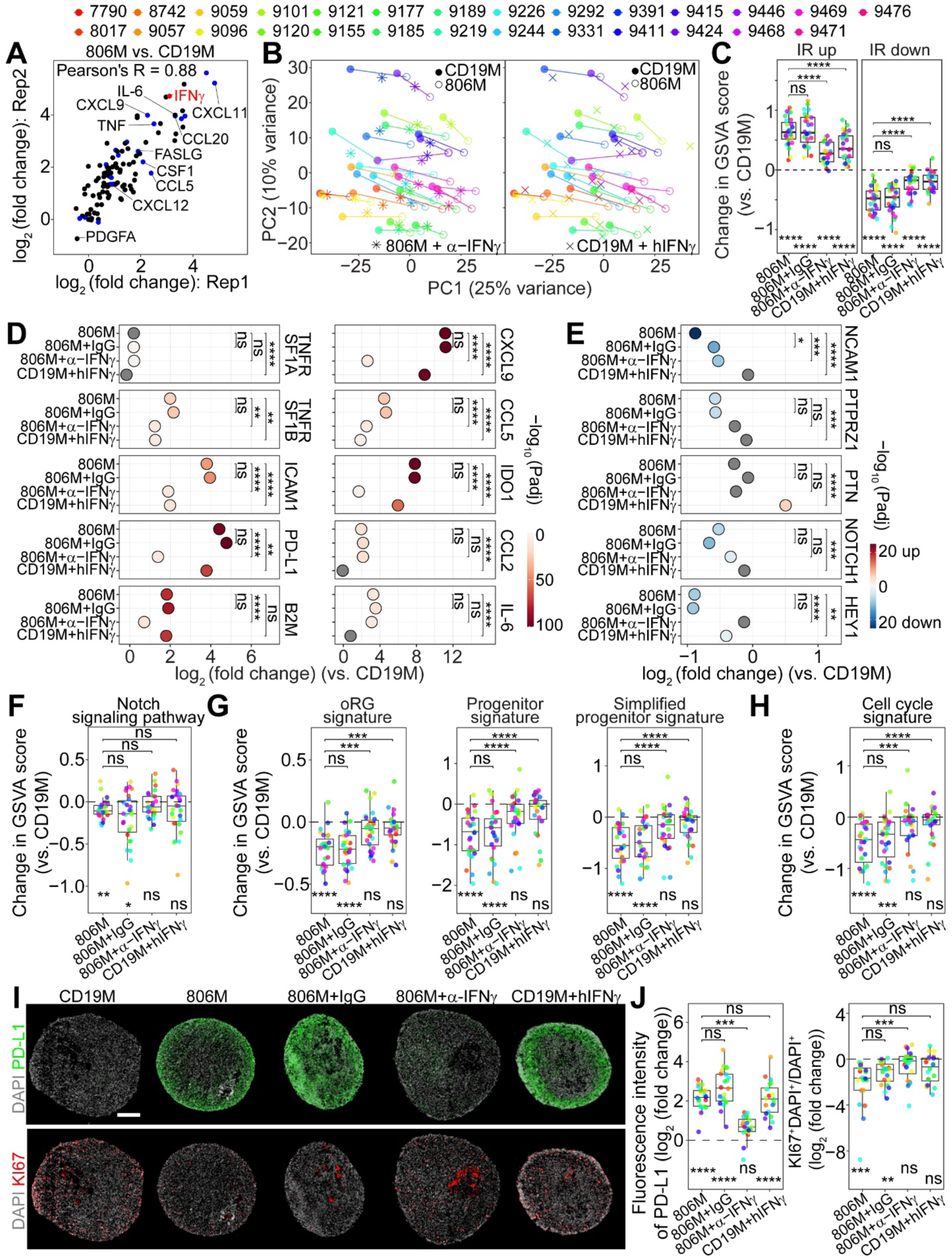
IFNɣ is required but not sufficient to mediate GBO and CAR-T cell co-culture conditioned medium-induced downregulation of glioma stem cell signatures and reduced cell proliferation. (A) Scatterplot of cytokine array results as log_2_fold-change in 806M vs. CD19M across two biological replicates. IFNɣ is highlighted as one of the top upregulated proteins (red dot). Other upregulated cytokines also identified by scRNA-seq are highlighted in blue. (B) PCA plots of IFNɣ depletion in 806M (with 10 ng/mL anti-IFNɣ neutralizing antibody; left) and IFNɣ (10 ng/mL) repletion in CD19M (right) with reference points of CD19M and 806M alone (same as in Figure 6A). Note that IFNɣ depletion in 806M led to an overall partial left-shift back towards CD19M, and IFNɣ repletion in CD19M led to an overall partial right-shift forwards towards 806M for all samples. (C) Boxplot and jitterplot of change in GSVA score compared to CD19M of up- and down-regulated IR signatures from scRNA-seq dataset as in Figure 2C. Asterisks below the boxplots and without brackets indicate statistical comparisons with CD19M. Asterisks with brackets correspond to pair-wise statistical comparisons as indicated (ns: p ≥ 0.05; *p < 0.05; **p < 1e2; ***p < 1e3;****p < 1e4; paired student’s t-test with Bonferonni correction). (D) Scatterplot of upregulated ligand and receptor molecules associated with immune inhibitory interactions and leukocyte recruitment. X-axis indicates the log_2_fold-change of gene expression versus CD19M, and point color indicates the -log_10_P-value of the differential expression versus CD19M. Asterisks with brackets correspond to the statistical comparisons as indicated. (ns: p ≥ 0.05; *p < 0.05; **p < 1e2; ***p < 1e3;****p < 1e4; DESeq2 with Benjamini-Hochburg correction). (E) Scatterplot of ligand and receptor molecules associated with downregulated cell-cell interactions within the neoplastic cell populations. Plot is formatted as in (D) (F-H) Boxplot and jitterplot of change in GSVA score from CD19M of Notch pathway (F), glioma stem cell (G), and cell cycle (H) signatures. Statistics are formatted as in (C) (I) Sample confocal images of immunostaining images for PD-L1 and KI67 in UP-9185 under different conditions. Scale bar: 200 μm. (J) Quantification of PD-L1 immunostaining intensity and percentage of KI67^+^ cells in GBOs derived from 20 patient samples shown as log_2_fold-change from the CD19M condition. Statistics are formatted as in (C) Also see **Figure S7** and **Tables S5** and **S7**.

To directly examine the role of IFNɣ, we depleted IFNɣ from 806M with a neutralizing antibody and repleted IFNɣ levels in CD19M to those found in 806M with recombinant human IFNɣ proteins (See Star Methods). We performed bulk RNA-seq of GBOs from the same 27 patient samples 3 days after treatment. PCA analysis showed that overall IFNɣ depletion in 806M led to a partial shift leftwards towards CD19M, whereas IFNɣ repletion in CD19M led to a partial shift rightwards towards 806M (Figure 7B). GBOs exposed to 806M with an isotype control IgG antibody exhibited few changes compared to 806M (Figure S7B-C). The IR signature was weakened upon IFNɣ depletion in 806M, though IFNɣ repletion in CD19M was inadequate to fully recapitulate the effect of 806M (Figure 7C). Analysis of individual genes showed that some changes were IFNɣ-dependent, such as PD-L1, IDO1, CXCL9 and MHC I component B2M, whereas others were IFNɣ-independent, such as CCL2 and IL-6 (Figure 7D).

For genes involved in cell-cell interactions among neoplastic cells, IFNɣ also showed complex effects (Figure 7E). Downregulation of PTPRZ and Notch signaling, as well as glioma stem cell and cell cycle signatures, was abolished by depletion of IFNɣ in 806M, but IFNɣ repletion in CD19M failed to fully recapitulate the phenotype (Figure 7E-H). This trend was also observed at the individual gene level, including cell cycle genes MKI67 and TOP2A, and oRG-related genes, FAM107A and NESTIN (NES) (Figure S7C-D). Immunostaining of PD-L1 confirmed that IFNɣ depletion in 806M led to a significant downregulation, whereas IFNɣ repletion in CD19M restored protein expression to similar levels as 806M (Figure 7I-J and S7E). Cell proliferation as marked by Ki67 was restored upon depletion of IFNɣ in 806M, but repletion of IFNɣ in CD19M had no effect (Figure 7I-J and S7E).

Together, these results suggest that IFNɣ is required, but not sufficient, to fully recapitulate CAR-T cell treatment-induced downregulation of glioma stem cell signatures and reduced tumor cell proliferation.

## DISCUSSION

CAR-T cell therapy has achieved remarkable outcomes in several difficult-to-treat cancers, however, its application in GBM faces unique and poorly understood challenges that have thus far limited its success. To gain insight into these challenges, we performed longitudinal, multi-modal, single-cell analyses of a patient-derived glioblastoma organoid model of CAR-T cell therapy. An adaptive tumor response is initiated by CAR-T cell activity, which leads to extensive shifts in gene expression and changes in cell-cell interaction networks with both anti-tumor and pro-tumor properties, ultimately resulting in tumor persistence with T cell exhaustion. Our study revealed the initial anti-tumor responses and the extent of the later immune suppressive adaptive responses, which involves all cell types, and our single-cell analyses further define the contribution of various cell types within the tumor to specific signaling pathways. Surprisingly, the neoplastic cell response is also accompanied by the attenuation of glioma stem-like states and reduced cell proliferation across both antigen^+^ and antigen^-^ neoplastic cells, which can be recapitulated with conditioned media and requires IFNɣ. These findings indicate that soluble factors produced by immune activity within GBM, such as IFNɣ, can favorably alter neoplastic cell states, suggesting the potential of CAR-T cell therapy to broadly affect neoplastic cells, including antigen^-^ neoplastic cells, to improve overall treatment efficacy.

### A Single-cell Resource of Dynamic Gene Expression among Different Cell Types upon CAR-T Cell Treatment of GBM

The cellular heterogeneity and native cell-cell interactions in GBOs paired with temporal, multi-modal, single-cell analyses provide a powerful system for dissecting the complexity of CAR-T cell treatment responses to understand how dynamic interactions among different cell types drive broad biological phenotypes. The inclusion of multiple CAR constructs and GBOs derived from multiple patients with varying EGFRvIII levels demonstrate the replicability and relative effect sizes of our findings across this heterogeneous disease. We further confirmed the major findings from single-cell analyses using paired patient samples before and shortly after CAR-T cell treatment in a clinical trial as well as GBOs derived from a large cohort of patients with GBM that contained diverse somatic mutations.

Given the difficulties in obtaining patient samples at multiple time points for dynamic analysis, this rich dataset will be a useful resource for the field and has significant implications for future treatment strategies. We found opposing tumor responses with some favorable anti-tumor effects, such as those promoting T cell recruitment and engraftment and production of cytotoxic factors, but also many more immunosuppressive changes at both the molecular and cellular levels. Previous observations of the upregulation of some inhibitory immune checkpoint molecules, such as PD-1, have prompted the exploration of combination therapies of CAR-T cells with anti-PD-1 blockade in clinical trials (NCT03726515, NCT04003649); however, this approach may yield limited benefits as other untargeted pathways remain active in exerting immune suppressive activity^53^, which could help to explain a lack of efficacy in a recent clinical trial for GBM^89^. We also observed the production of several different recruitment and differentiation signals that promote the accumulation of inhibitory immune regulatory cells, such as Tregs, M2-like macrophages, and MDSCs, suggesting that the desired effects of targeted agents, such as those antagonizing CCL2 or CSF1 signaling, may be circumvented by other pathways or cell types^61,90^. It may be possible nonetheless that intervention in a clinically manageable subset of these mechanisms may yield a therapeutically meaningful outcome, and the prioritization and assessment of these approaches should be an area of future investigation. Importantly, we found upregulation of many chemokines that can recruit T cells, which provides mechanistic insight into the major finding from the previous 2173 clinical trial, namely that peripherally infused CAR-T cells can traffic to and infiltrate the tumor^26^. We also discovered several intrinsic pro-immune signals, including IL-15/IL15RA and OSM, which may be further enhanced to improve the immune activity and treatment responses.

### Targeting the “Non-targetable” with CAR-T Cells

Our most surprising finding is that soluble factors produced upon CAR-T cell treatment attenuated glioma stem-like states and reduced cell proliferation of both antigen^+^ and antigen^-^ neoplastic cells. In addition to the pursuit of sustained and enhanced tumor cell killing by CAR-T cells, our finding suggests that CAR-T therapy can have a direct cytostatic effect on neoplastic cells that are negative for the target antigen, thus broadening the potential scope of therapeutic benefit. While our study focused on CAR-T cells against EGFRvIII, this finding is likely to be applicable to CAR-T cell treatment for other targets, including in the recent phase 1 trial of bivalent CAR-T targeted against both IL13RA2 and EGFR^7^. Additionally, a prominent case report of IL13RA2-targeted CAR-T therapy in recurrent multifocal GBM demonstrated remarkable tumor regression despite the highly variable expression of the target antigen^91^. Although it was hypothesized that the host immune system was recruited and engaged by CAR-T cell activity to further promote antitumor responses, soluble factors generated upon immune activity might have also contributed by acting directly on neoplastic cells in a favorable manner. IFNɣ is a critical mediator at the crossroads of these interactions with opposing functions coordinating both immune-suppressive and activating responses^92^. In GBM specifically, a recent study in mice suggested that IFNɣ plays a critical role in recruiting endogenous T cell and monocytes/macrophages, both of which play a substantial role in the overall treatment response^56^. Identification of the factors involved in regulating these functions, as well as their temporal context, may reveal opportunities to further tilt the balance in favor of enhancing anti-tumor activity.

While there are several models for cell hierarchies in GBM, many agree on the presence of adaptive cellular plasticity that mediates treatment resistance and is enriched in glioma stem-like states^14^. These glioma stem cells are proposed to exhibit signaling promiscuity and epigenetic deregulation, which enable them to generate the observed cellular heterogeneity in GBM and adapt to therapeutic challenges. We found that soluble factors produced upon CAR-T cell treatment in GBOs attenuated glioma stem-like states and reduced cell proliferation for GBOs derived from a large cohort of patients with diverse somatic mutation landscapes and established glioma stem cell lines from multiple patients. We further identified IFNɣ as one of the diffusible factors required, but not sufficient, for such an effect. We also observed a correlation between the initial EGFRvIII levels and the degree of downregulation of key stem cell factors (Figure 2C-D), suggesting that this response may be dose-dependent and that enhanced production of these soluble factors would broaden clinical translation.

Our findings support an additional role for CAR-T cell therapy in GBM as a delivery mechanism of potent soluble factors that attenuate glioma stem cell activity and reduce proliferation of both antigen^+^ and antigen^-^ neoplastic cells. Together with the observed multi-factorial immune suppressive adaptive tumor response, these findings point towards maximizing the efficacy of an early and limited window of opportunity with complementary approaches that augment CAR-T cell activity. Such an approach may include increasing initial on-target activity by construction of a CAR-T that targets multiple antigens, as evidenced by the comparative success of a recent trial evaluating a dual-EGFR/IL13RA2 CAR-T^7^. Additionally, CAR-T cells could potentially be engineered with diffusible payloads that exert cytostatic and cell state-modulating activity. These anti-tumor effects can persist even after T cell exhaustion, suggesting that ongoing immune activity may not be necessarily required for beneficial results. Cytokines have wide ranging, pleiotropic, and cell type-specific effects, as demonstrated by the growing body of work studying IFNɣ in immunotherapy^93^, and ongoing efforts to engineer synthetic variants with greater selectivity for specific receptor or receptor complexes involved in desired functions may enable their clinical use.

### Limitations of the Current Study

First, there exists tremendous heterogeneity in GBM, and we only focused on GBOs from five patients for detailed temporal single-cell analyses in response to three different CAR-T types. Future studies with samples from more patients with different somatic alterations may confirm these findings. So far, we have validated our major findings with GBOs from additional twenty-seven patient samples with diverse somatic mutation landscapes (Table S1). Second, for direct comparison to gain mechanistic insight, we used the same three CAR-T cells for different GBOs. Given the potential native T cell dysfunction in GBM^94^, future studies can apply paired patient-derived CAR-T cells and GBOs for analysis as in our recent study (see accompanying manuscript). Third, there are limitations of our *in vitro* GBO and CAR-T cell co-culture model. For example, the model currently lacks a peripheral immune system and does not permit the direct examination of the recruitment of immune cells from the circulatory system. We validated the key findings using patient-derived glioma stem cell lines and rare clinical samples that were paired before and shortly after CAR-T cell treatment from the same patients, but the number of patients was limited and the timing of tissue sampling was not well controlled. Future studies with more patient samples from additional clinical trials will validate these findings. Fourth, we identified IFNɣ as a diffusible factor required but not sufficient to reduce glioma stem cell-like signatures and proliferation of both antigen^+^ and antigen^-^ neoplastic cells upon CAR-T cell treatment. Additional diffusible factors remain to be identified.

In summary, our single-cell analyses of an organoid model of CAR-T cell therapy in GBM revealed the dynamic evolution of tumor adaptive responses and uncovered an opportunity to leverage soluble factors produced with immune activity to broadly target neoplastic cells.

## METHODS

### EXPERIMENTAL MODEL AND SUBJECT DETAILS

#### Human Subjects

The use of brain tumor tissue was coordinated and overseen by the University of Pennsylvania Tumor Tissue/Biospecimen Bank following ethical and technical guidelines for the use of human samples in biomedical research. Patient GBM tissues were collected at the Hospital of the University of Pennsylvania after informed patient consent under a protocol approved by the Institutional Review Board of University of Pennsylvania. All patient samples were de-identified prior receipt and processing. A total of 36 patient cases were included in the present study, including 5 for single-cell analyses, 5 from a previous clinical trial (NCT02209376) with tumor samples before and shortly after 2173 CAR-T cell treatment, and 27 for conditioned media experiments. Table S1 lists the epidemiological data for each subject and genomic data for each tumor provided by the Neurosurgery Clinical Research Division (NCRD) at the University of Pennsylvania and the University of Pennsylvania Center for Personalized Diagnostics. Genetic testing was performed using a targeted panel of disease-associated genomic alterations (Agilent Haloplex assay, Illumina HiSeq2500) and fusion transcripts (Illumina HiSeq2500). MGMT promoter methylation was analyzed using PyroMark Q24 (Qiagen).

### Generation and Culture of Glioblastoma Organoids (GBOs) from Patient Samples

GBOs were generated and cultured using our established protocol as previously described^22,23^. Briefly, surgically resected GBM tissue was transported to the laboratory in Hibernate A (Thermo Fisher Scientific) and dissected in Hibernate A with 1X GlutaMAX (Thermo Fisher Scientific) and 1X Antibiotic-Antimycotic (Thermo Fisher Scientific) into small 0.5-1.0 mm in diameter pieces using small spring scissors (Fine Science Tools) in a sterile Petri dish (VWR). Tumor pieces were treated with red blood cell lysis buffer (Thermo Fisher Scientific) and washed with GBO medium containing 50% DMEM:F12 (Thermo Fisher Scientific), 50% Neurobasal (Thermo Fisher Scientific), 1X GlutaMax, 1X NEAAs (Thermo Fisher Scientific), 50 U/mL Penicillin-Streptomycin (Thermo Fisher Scientific), 1X N2 supplement (Thermo Fisher Scientific), 1X B27 w/o vitamin A supplement (Thermo Fisher Scientific), 1X 2-mercaptoethanol (Thermo Fisher Scientific), and 2.5 μg/ml human insulin (Sigma). Samples were cultured in 4 mL GBO medium in a 6-well flat-bottom culture plate (Corning) on an orbital shaker equipped with a rubber mat platform (Thermo Fisher Scientific). Media were changed every 2 days, and GBOs were passaged by microdissection when grown to sizes ∼ 1.0 mm in diameter. UP-7790 and UP-8017 were retrieved from our biobank^22,23^ by warming in a 37 °C water bath until thawed, washed after dropwise addition of 10 mL GBO medium supplemented with 10 µM Y-27632 (StemCell Technologies), and cultured for one day in the presence of 10 µM Y-27632.

### Co-culture of GBOs with CAR-T Cells

CAR-T cells were generated in parallel from the same healthy donor for all three CARs as previously described (28), and frozen in aliquots of 1×10^6^ cells/vial. Cryopreserved CAR-T cells were warmed in a 37 °C water bath until thawed, washed after dropwise addition of 10 mL T cell medium, containing ImmunoCult™-XF T Cell Expansion Medium (StemCell Technologies) and 50 U/mL Penicillin-Streptomycin (Thermo Fisher Scientific), and cultured for 1 day in the presence of 100 units of IL-2 (StemCell Technologies) followed by 2 days in T cell medium alone. The number of cells per GBO was determined empirically by dissociation with the Brain Tumor Dissociation Kit (Miltenyi), and was generally in the range of 100-500K per GBO. GBOs and CAR-T cells were co-cultured at a 10:1 starting ratio of cells, and care was taken to ensure that each well contained a similar number of GBOs. Co-culture was performed in T cell medium for the first day and the medium was completely removed the next day, including non-engrafted T cells in the medium, and then cultured in GBO medium followed by media changes every 2 days (Figure S1A). The current approach better mimicked the clinically observed overall treatment resistance (Figure 1C). Live imaging of CAR-T cell and GBO co-culture was performed with a combination of CellTracker dyes (ThermoFisher Scientific) pre-incubated with either 9057 GBOs or 806 CAR-T cells and a caspase-3/7 dye, imaged with a Zeiss LSM 710 confocal.

### IFNɣ ELISA

For IFNɣ ELISA analysis, 50 μL of culture medium was sampled, snap frozen on dry ice, and stored at -80°C. Individual wells of GBO and CAR-T cell co-cultures were considered as biological replicates, and an ELISA was performed with three biological replicates on 10-fold diluted medium using human IFNɣ DuoSet ELISA kits (R&D Systems). The absorbance was recorded at 450 nm using a microplate absorbance reader (Tecan) with a wavelength correction set at 540 nm. The signal from the blank condition was subtracted from all the standards and experimental values. A 7-point standard curve was derived from a linear fit of the log_10_ transformed IFNɣ concentration versus the log_10_ transformed absorbance value of the standards and was used to calculate sample IFNɣ concentrations.

### Tissue Processing, Immunohistology, and Confocal Imaging

For histological analysis, patient tissues or GBOs were fixed with 4% methanol-free formaldehyde (Polysciences) in DPBS (Thermo Fisher Scientific) for 30 min at room temperature with rotation (VWR), followed by overnight incubation in 30% sucrose (VWR) in DPBS at 4°C. GBOs were transferred to a plastic cryomold (Electron Microscopy Sciences), frozen in tissue freezing medium (Thermo Fisher Scientific), and stored at -80°C.

The samples were sectioned at 20 µm using a cryostat (Leica, CM3050S) and transferred onto charged slides (Thermo Fisher Scientific). The slides were dried at room temperature and stored at - 20°C. For immunostaining, the tissue sections were outlined with a hydrophobic pen (Vector Laboratories), transferred to a moisture chamber (Newcomer Supply), and washed 1x with TBST (Thermo Fisher Scientific). Tissue sections were permeabilized and blocked using a solution containing 10% donkey serum (MilliporeSigma), 0.5% Triton X-100 (MilliporeSigma), and 22.52 mg/ml glycine (MilliporeSigma) in TBST for 1 hr at room temperature. Tissue sections were incubated with primary antibodies diluted in TBST with 5% donkey serum and 0.1% Triton X-100 overnight at 4°C. After washing 3x in TBST, tissue sections were incubated with secondary antibodies diluted in TBST with 5% donkey serum and 0.1% Triton X-100 for 1 hr at room temperature. After washing 3x with TBST, mounting medium (Vector Laboratories) was added and slides were cover-slipped (Fisher Scientific). For the patient samples, an additional autofluorescence quenching step was performed prior to mounting. The slides were washed 1X with DPBS, treated with 1X TrueBlack (Biotium) diluted according to the manufacturer’s instructions for 30 s, and then washed 3X with DPBS.

Histology slides were imaged as z-stacks with a Zeiss LSM 810 confocal microscope or a Zeiss LSM 710 confocal microscope (Zeiss) using a 20X objective with Zen 2 software (Zeiss). Tiled images were taken to capture the entire organoid to better reveal any intra-organoid heterogeneity and to represent any observed phenotype more completely. Typically, GBO sections can be fully captured with 3 x 3 tiles and 5 z-stacks of 1.6 µm. Images were pre-processed by stitching and orthogonal projection using Zen 2. Custom ImageJ analysis pipelines were used for the quantification of immunostaining images. Importantly, sets of images that were compared with each other (e.g., between different conditions of GBOs derived from the same patient) were generated in parallel with the same immunostaining solutions and microscopy settings.

### Multi-modal Single-cell Analyses Library Construction

Co-cultures using GBOs from the same patient tumor but with different CAR-T cells were performed, processed, and analyzed in parallel. Approximately 50 GBOs collected at each time point were dissociated using the Brain Tumor Dissociation kit (Miltenyi) with the following modifications to the manufacturer’s protocol. Mechanical tissue disruption was performed by manual pipetting for 10-20 strokes three times: at the start of enzyme incubation, after 15 min, and after an additional 10 min. A P1000 tip was used for the first time, a P200 tip nested over a P1000 tip was used for the second time, and a P10 tip nested over a P1000 tip was used for the third time. Additional mechanical disruption focused on the remaining tissue chunks at either or both the second and third time point by allowing the tissue chunks to settle under gravity, removing the tissue chunks, and pipetting up and down using either a P200 or P10 pipette tip in a separate 1.5 mL Eppendorf tube. We found that this protocol minimized mechanical damage to the already dissociated cells while allowing for more complete GBO dissociation. All incubations were performed at 37 °C on a tube rotator. To minimize temperature changes during the mechanical disruption steps, especially while processing multiple samples, samples were kept at 37 °C when removed from the incubator. The crude cell suspension was filtered through a 70 µm mesh (Miltenyi), diluted in 10 mL DPBS containing 2% BSA (VWR), spun down at 250 g for 5 min at 4 °C, and resuspended in 250 µL ice cold CITE-seq staining buffer, containing 2% BSA and 0.01% Tween (Sigma) in DPBS. A small aliquot was used for trypan blue staining and automated cell counting to ensure viability > 85% and yield > 0.5 million cells.

CITE-seq staining, droplet encapsulation on our drop-seq platform, and library preparation were performed as previously described with some modifications^22,31^. A pool of 15 barcoded antibodies (BioLegend; Table S2) was used and antibody staining was performed at 4 °C with gentle rotation. After PCR amplification, the PCR tube was briefly centrifuged, and 48 µL of the reaction mixture was retrieved from each tube and pooled for purification. The final ADT libraries were amplified with 8 cycles. Both scRNA-seq and ADT libraries were analyzed using a bioanalyzer and quantified by qPCR (KAPA) with appropriate corrections for mean library size. Samples were pooled, loaded at 3.0 pM, and sequenced on a NextSeq 550 (Illumina) with 20 bp Read 1, 8 bp Index 1, and 64 bp Read 2. The Drop-seq custom Read 1 primer was spiked into the Illumina sequencing primer well (#20) at the manufacturer’s recommended concentration.

### EGFRvIII Genotyping Library Construction and Data Preprocessing

EGFRvIII genotyping from full-length scRNA-seq cDNA was performed using an optimized method based on the published Gibson cloning-based Genotyping of Transcriptomes protocol^33^ (Figure S1G). The two modifications included: 1) use of an oligo-dA primer for PCR #2 as there was substantial heterogeneity in the coverage at the 3’ end of the transcript; and 2) use of two primers in PCR #3 targeting exons 2 and 8 to yield fragments that were of similar length in the final sequencing library.

PCR #1 was set up with 10 ng amplified cDNA in 10.5 µL, 1 µL of 10 µM PCR1_fw primer, 1 µL of 10 µM PCR1_rv primer, and 12.5 µL of 2X KAPA HiFi HotStart ReadyMix (Roche). The thermocycling conditions were as follows: 98 °C for 3 min; 12 cycles of 98 °C for 20 s, 65 °C for 30 s, and 72 °C for 2 min; and finally 72 °C for 5 min. The PCR product was purified with 1.0X Ampure XP beads (Roche) and eluted in 10 µL H_2_O. The final DNA concentrations were measured using the Qubit dsDNA HS Assay Kit (Thermo Fisher Scientific).

Circularization #1 was set up with 890 µL H_2_O, 100 µL 10X CutSmart Buffer (NEB), ∼200 ng PCR #1 product in 10 µL, and 10 µL of Gibson Assembly Master Mix (NEB). After incubation at 50 °C for 1 h, DNA was purified with 1.0X Ampure XP beads and eluted in 10 µL H_2_O.

PCR #2 was set up with 50 ng DNA in 10.5 µL, 1 µL of 10 µM PCR2_fw primer, 0.5 µL of 1 µM PCR2_rv_exon2 primer, 0.5 µL of 1 µM PCR2_rv_exon8 primer, and 12.5 µL of 2X KAPA HiFi HotStart ReadyMix. The thermocycling conditions were as follows: 98 °C for 3 min; 12 cycles of 98 °C for 20 s, 46 °C for 30 s, and 72 °C for 2 min; and finally 72 °C for 5 min. PCR product was purified with 1.0X Ampure XP beads and eluted in 10 µL H_2_O. Final DNA concentrations were measured using the Qubit dsDNA HS Assay Kit. In optimization experiments, PCR2_fw_UTRstart, which was complementary to the start of the 3’ UTR, and PCR2_fw_UTRend, which was complementary to the end of the 3’ UTR, were tested.

Circularization #2 was set up with 890 µL H_2_O, 100 µL 10X CutSmart Buffer (NEB), ∼25 ng PCR #2 product in 10 µL, and 10 µL of Gibson Assembly Master Mix. After incubation at 50 °C for 1 h, DNA was purified with 1.0X Ampure XP beads and eluted in 10 µL H_2_O.

PCR #3 was set up with 10 ng DNA in 10.5 µL, 1 µL of 10 µM PCR3_fw primer, 1 µL of 10 µM P5-TSO primer, and 12.5 µL of 2X KAPA HiFi HotStart ReadyMix. Thermocycling conditions were as follows: 98 °C for 3 min; 8 cycles of 98 °C for 20 s, 65 °C for 30 s, and 72 °C for 30 s; and finally 72 °C for 5 min. The PCR product was purified with 1.0X Ampure XP beads and eluted in 10 µL H_2_O.

PCR #4 was set up with 2 ng DNA in 10.5 µL, 1 µL of 10 µM RPI-X primer (Illumina), 1 µL of 10 µM P5-TSO primer, and 12.5 µL of 2X KAPA HiFi HotStart ReadyMix. Thermocycling conditions were as follows: 98 °C for 3 min; 6 cycles of 98 °C for 20 s, 65 °C for 30 s, and 72 °C for 30 s; and finally 72 °C for 5 min. The PCR product was purified with 1.0X Ampure XP beads and eluted in 10 µL H_2_O.

Final GoT libraries were analyzed using a bioanalyzer (Agilent) with expected short fragments (exon1-2 of EGFRvIII^-^ transcripts and exon1-8 of EGFRvIII^+^ transcripts) at ∼260 bp and a long fragment (exon1-8 of EGFRvIII^-^ transcripts) at ∼1,100 bp. Additional small peaks were observed between these short and long fragments, which appeared to correspond to low abundance splice variants within the long first intron of EGFR. GoT libraries were quantified by qPCR, pooled with the scRNA-seq and ADT libraries, and sequenced as described above.

Linear genotyping was performed by targeted enrichment and PCR^96^. PCR #1 was set up with 10 ng amplified cDNA, 1.25 µL of 6 µM SMART-AC primer, 1.25 µL of 6 µM linear_PCR1_fw primer, and 12.5 µL of 2X KAPA HiFi HotStart ReadyMix. Thermocycling conditions were as follows: 95 °C for 3 min; 10 cycles of 98 °C for 20 s, 65 °C for 15 s, and 72 °C for 3 min; and finally 72 °C for 5 min. The PCR products were purified with 0.7X Ampure XP beads and eluted in 20 µL H_2_O. Biotinylated fragments were captured using magnetic Dynabeads kilobaseBINDER streptavidin beads (Thermo Fisher Scientific) according to the manufacturer’s protocol. PCR #2 was set up with 23 µL bead suspension in H_2_O, 1 µL of 10 µM Nexterra N7XX (Illumina) primer, 1 µL of 10 µM P5-TSO, and 25 µL PfuUltra II Hotstart 2x Master Mix (Agilent). Thermocycling conditions were as follows: 95 °C for 2 min; 4 cycles of 95 °C for 20 s, 65 °C for 20 s, and 72 °C for 2 min; 6 cycles of 95 °C for 20 s, 72 °C for 2 min and 20 s; and finally 72 °C for 5 minutes. Streptavidin beads were magnetized, and the supernatant was collected for 0.7X Ampure XP bead purification. The final library was eluted in 10 µL H_2_O, analyzed by bioanalyzer, quantified by qPCR, and pooled with the scRNA-seq libraries, ADT libraries, and GoT based EGFRvIII genotyping libraries for sequencing under the same conditions.

### Single-cell Multi-modal Data Pre-processing

Raw sequencing data were demultiplexed using bcl2fastq2 (Illumina) with adapter trimming turned off. Additional processing was performed using drop-seq tools v.2.1.0 with GRCh38 as the reference genome and gencode v.28 GTF as the annotation file. The gene expression matrix was retrieved from reads that overlapped exclusively with exons or UTRs. Barcodes associated with ambient RNA were identified using EmptyDrops^97^ and removed from the analysis. Barcodes associated with doublets were identified using Scrublet^98^ and were removed. Barcode clashes between samples were also removed from the dataset. Barcodes corresponding to ∼122K individual cells were included in the final gene expression matrix.

The whitelist of the barcodes was used to retrieve ADT counts using CITE-seq-Counts^99^ with cell barcodes and UMI settings adjusted for Drop-seq (-cbf 1 -cbl 12 -umif 13 -umil 20). This whitelist was also used to retrieve EGFRvIII genotyping data using IronThrone-GoT^33^ with cell barcodes and UMI settings adjusted for Drop-seq (--bclen 12 --umilen 8). A custom config file was provided with a common exon1 sequence (GGCTCTGGAGGAAAAGAAAG) at base pair positions 1-20, an exon 2 sequence (TTTGCCAAGGCACGAGTAACAAGCTC) assigned to EGFRvIII wild-type at base pair positions 21-46, and exon 8 sequence (GTAATTATGTGGTGACAGATCACGGC) assigned to EGFRvIII mutant at base pair positions 21-46. UMIs with fewer than 10 reads or with both wild-type and mutant reads were excluded. Cells containing EGFRvIII wild-type only UMIs were called as EGFRvIII^-^, whereas cells containing any EGFRvIII mutant UMIs were called as EGFRvIII^+^. In the EGFRvIII optimization experiments, sequencing data were processed in the same way for both GoT based and linear approaches. Alignment was performed using STAR and on-target alignments uniquely corresponding to exon1 were quantified using bedtools^100^. Alignment coverage was visualized using IGV^101^.

### Single-cell Data Normalization and Clustering

Gene expression, ADT, and EGFRvIII genotyping data were integrated into a single object using Seurat v3.2^102^. The scRNA-seq counts were normalized using SCTransform^103^, and coarse clustering and UMAP dimensionality reduction were performed using the top 75 principal components. ADT counts were normalized using centered log ratio transform.

Neoplastic cells were identified based on the presence of large chromosomal aberrations. As EGFR mutant GBMs commonly demonstrate chromosome 7 amplification and chromosome 10 deletion^32^, we performed enrichment analysis for genes based on chromosomal position using the AddModuleScore function and calculated the ratio between the corresponding enrichments. An additional copy number variation analysis was performed using inferCNV (inferCNV of the Trinity CTAT Project, https://github.com/broadinstitute/inferCNV). For computational feasibility, we analyzed each patient sample individually with macrophages/microglia and T-cells set as the normal reference. Non-neoplastic cell subtypes were identified by the presence of marker genes.

Neoplastic cells in GBOs from different patients were further subclustered to identify basal versus immune responsive (IR) populations. Non-neoplastic cells and T cells exhibited variable abundance in different samples and were thus pooled together by cell type for further subclustering to identify different states. T cell gene expression was further denoised using SAVER-X, which has been consistently found to be a reliable method in benchmarking studies^104^. As SAVER-X denoising yields a dense matrix with substantially fewer zeros, marker gene expression was shown using the kernel density function in Nebulosa^105^.

### Gene Signature Identification and Enrichment Analysis

Differential gene expression analysis between the basal vs. IR populations was performed using the FindMarkers function in Seurat. Gene ontology and transcription factor enrichment analysis was performed using gProfiler2^95^ using differentially expressed genes (DEGs) with an adjusted P value < 0.05 and ranked by log transformed fold-change.

Enrichment scores for specific pathways were calculated using the AddModuleScore function. The molecular signatures of different pathways and cell types used for the analysis are listed in Table S4. For the PTPRZ1 KD signature, since both up- and downregulated genes were available^83^, the overall enrichment score was taken as the difference between the individual enrichment scores of the upregulated genes and the enrichment of the downregulated genes. Cell cycle assignment was determined using the CellCycleScoring function in Seurat v.3.2. For separate analysis of EGFRvIII^+^ and EGFRvIII^-^ neoplastic cells, only cells from experiments with 2173 were included in the analysis as this 2173 is specific to EGFRvIII, whereas 806 has reactivity against wild-type EGFR and other mutants.

### Ligand-receptor Interaction Analysis

Ligand-receptor analysis was performed using the database and statistical methods implemented in CellPhoneDB as previously described^39^. The aggregated gene expression dataset was used as input and statistically significant interactions were defined as P < 0.05. The number of significant interactions was quantified using the heatmap_plot function, and chord diagram visualization was performed using Circlize^106^. The expression of the ligand and receptor was included in the output of CellPhoneDB, and it was taken as a measure of the interaction strength as previously described^39^. Interactions that were not statistically significant were set to a strength of zero.

To compare interactions between different states (e.g., basal vs. IR, T active vs. T start, etc.), the ratio of the strengths of each individual interaction was calculated. Histograms of log_10_ transformed results were fitted with a kernel density function, and interactions that were only significant in one of the two comparator states were arbitrarily assigned a value of ± 1.5 for visualization purposes (Figure 3C and S3A-B). For gene ontology enrichment analysis with gprofiler2, individual genes corresponding to interactions with a greater than 1.25-fold change in strength between the two comparator conditions were used as an unranked list. Importantly, background genes were set to all genes included in the CellPhoneDB database.

To analyze the interactions among neoplastic cells (Figure 5 and S5), neoplastic cells were subclustered and GBOs from different patients were analyzed separately as described above. To examine how the interactions changed from basal to IR states, the subclusters were marked as belonging largely to either the basal or IR state. The numbers of significant interactions within different basal clusters, between basal and IR clusters, and within different IR clusters were quantified. To account for the different number of clusters in the different groups, the number of significant interactions was divided by the total number of possible interactions for each category to obtain the percentage of significant interactions. To evaluate monotonicity, we took these three groups as ordinal variables in the order listed above, and Spearman’s rho was calculated against the percentage of significant interactions for each individual ligand-receptor pair. To account for the degree of change, the calculated Spearman’s rho was multiplied by the absolute value of the difference in percent significant interactions between the basal only and IR only interactions, yielding the IR vs. basal enrichment score (Figure 5A).

### Conditioned Media, Cytokine Array, and IFNɣ Experiments

Conditioned media was collected from co-cultures of GBOs (8017, 9057, and 9121) with CAR-T cells (CD19, 2173, and 806) at days 2, 3, and 4. This time window was selected because it demonstrated peak T cell activity, as determined by IFNɣ production, and contained the GBO media, not the T cell media. Media from different GBOs and days were pooled and centrifuged at 500 g for 5 min to remove cells and debris. The supernatant was filtered through a 0.2 µm filter, aliquoted, and stored at -20 °C. This entire procedure was repeated twice to yield two biological replicates.

Cytokine array analysis was performed using the Proteome Profiler Human XL Cytokine Array Kit (R&D Systems), which includes 105 targets. The same 500 µL conditioned media as above was used as input, and the array was performed across both biological replicates. CD19 and 806 conditioned media membranes for each replicate were imaged simultaneously. Images were analyzed in ImageJ using the Protein Array Analyzer tool. Spot intensities were normalized to the positive and negative controls within each membrane and averaged between the two technical replicates. Spots with intensities greater than 5 standard deviations above that of negative controls were considered to be reliably detected and retained for further analysis (Table S7).

For activity validation and concentration determination, anti-IFNɣ neutralizing antibody (R&D Systems) was titrated into 806 conditioned media starting at 50 µg/mL with 10 serial 3-fold dilutions down to 0.85 ng/mL. A parallel titration of the mouse IgG_2A_ isotype control was performed to confirm non-reactivity. Antibody was added to 806 conditioned media, the mixture was incubated at room temperature for 30 min, and free IFNɣ levels were quantified by ELISA. A concentration of 10 ng/mL was selected as the anti-IFNɣ antibody showed near maximal neutralization and the isotype control showed no change in measured IFNɣ levels. Furthermore, levels of IFNɣ in the CD19 and 806 conditioned media were quantified to be 0 and ∼8 ng/mL, so human recombinant IFNɣ was added at a final concentration of 10 ng/mL in CD19 conditioned media for repletion experiments. Care was taken that IFNɣ was obtained from a non-bacterial source, in this case, from HEK293 cells, as post-translational modifications, such as glycosylation, could impact activity^107,108^.

For conditioned media experiments, aliquots of conditioned media were gently warmed to room temperature. Anti-IFNɣ antibody (10 ng/mL), mouse IgG_2A_ isotype control (10 ng/mL), and recombinant human IFNɣ (10 ng/mL) (all from R&D systems) were added to the appropriate conditioned media and incubated at room temperature for 30 min. GBOs were cultured in the conditioned media for 3 days before sample collection. Six GBOs were collected and analyzed by immunostaining, and three GBOs were pooled for a single bulk RNA-seq sample. In total, GBOs derived from 27 different patient samples were used for bulk-RNA-seq and 20 for immunostaining validation.

### Glioma Stem Cell Growth and Sphere Formation Assay

Glioma stem cell growth and sphere formation assays were performed as described using four previously established and characterized lines PBT003, PBT707, PBT726 and PBT111^86^. For cell growth, glioma stem cells were seeded at 5 x 10^4^ cells/well in 24-well plates and cultured in either CD19M or 806M supplemented with of 20 ng/mL epidermal growth factor (EGF) (PeproTech), 20 ng/mL fibroblast growth factor (FGF) (PeproTech). The cells were supplemented with fresh conditioned medium, EGF and FGF 3 days after culture. Cell growth was analyzed by cell counting 6 days after culture.

For sphere formation, glioma stem cells were seeded at 100 cells/well in 48-well plates and cultured in either CD19M or 806M supplemented with 20 ng/mL EGF, 20 ng/mL FGF for each conditioned medium. The cells were cultured with supplement of EGF and FGF every 3 days. Sphere formation was counted under a microscope 2 weeks after culture.

### RNA Isolation and Bulk RNA-seq Library Preparation

A collection of 5 µm thick FFPE sections from tumor samples before and shortly after CAR-T cell treatment of 5 patients who underwent the 2173 CAR-T cell clinical trial (NCT02209376)^26^ was obtained from the Division of Neuropathology at the University of Pennsylvania. RNA was isolated from a single section using the Quick-RNA FFPE Miniprep kit with on-column DNase I digestion (Zymo). RNA was analyzed by bioanalyzer and quantified using NanoDrop. Bulk RNA-seq libraries were constructed using the Illumina Stranded Total RNA Prep with the Ribo-Zero Plus kit (Illumina) according to the manufacturer’s instructions with 100 ng input, RNA fragmentation time reduced to 1 min, and 13 cycles of PCR amplification. Final libraries were additionally purified using 1.0X Ampure XP beads. Samples were pooled, loaded at 2.2 pM, and sequenced on a NextSeq 550 (Illumina) with 74 bp Read 1, 10 bp Index 1, 10 bp Index 2, and 74 bp Read 2.

For bulk RNA-seq analysis of GBOs, 3 organoids were pooled into a single sample, lysed in 400 µL TRIzol (Thermo Fisher Scientific), and stored at -80 °C. RNA isolation was performed according to the manufacturer’s instructions until partition of aqueous and organic layers. 200 µL of the aqueous layer was carefully retrieved and input into the RNA Clean and Concentrator Kit (Zymo) for further RNA purification. On-column DNase I digestion was performed according to the manufacturer’s instructions, and RNA was eluted in 15 µL of H_2_O. RNA was analyzed by bioanalyzer and quantified by nanodrop. Bulk RNA-seq libraries were constructed and sequenced as described above except that the full 2 min RNA fragmentation time was used.

### Bulk RNA-seq Analyses

Raw sequencing data were demultiplexed using bcl2fastq2 (Illumina) with adapter trimming turned off. Alignment was performed using STAR^109^, using the same reference genome as above, and gene expression was quantified using RSEM^110^. PCA of all samples was performed using the top 1000 most variable genes. Differential expression analysis was performed using DESeq2 with effect size estimation using Apeglm^111^. GO analysis was performed using gprofiler2 with DEGs having an adjusted P-value < 0.05 input as a list ranked by fold-change. Gene set enrichment was calculated using gsva ^112^ to yield the GSVA score. Comparison of GSVA scores for a given pathway between two conditions were performed using the Student’s paired t-test with Bonferroni correction for multiple hypothesis testing. The cell cycle signature was taken as the union of the S and G2/M phase genes used in the cell cycle analysis described above (Figure 5J and 6J).

### Quantification and Statistical Analysis

Differential gene and protein expression was determined in the scRNA-seq/CITE-seq analyses using the FindMarkers function in Seurat. A Wilcoxon rank-sum test was performed with Bonferroni correction for multiple hypothesis testing. Enrichment analyses for differentially expressed genes identified in the scRNA-seq data and for the genes involved in IR-enriched or basal-enriched interactions were performed using the statistical method implemented in gProfiler2 with the included g:SCS method for multiple hypothesis testing^113^. Notch pathway and stem cell signature enrichment analyses were performed using a paired Wilcoxon rank sum test. Statistical testing of the abundance of significant Notch interactions within neoplastic cell states was performed using one-way ANOVA (Figure S5G).

For the bulk RNA-seq analysis of 2173 trial patient samples and conditioned media experiments, differential gene expression was performed using DESeq2 with Benjamini-Hochburg correction for multiple hypothesis testing. GSVA scores for pathway enrichment for different conditions were compared using paired Student’s t-test. Unbiased enrichment analysis for the conditioned media experiments was performed using gProfiler2 as described for the scRNA-seq analysis above. Enrichment analysis for the conditioned media experiments using pathways identified earlier in the scRNA-seq experiments was performed using gene set enrichment analysis (GSEA) as implemented in fgsea with Benjamini-Hochburg correction for multiple hypothesis testing.

For confocal imaging analysis, nuclear staining (Ki67) quantification was performed as follows:

1) Zen 2 (.CZI) images for the DAPI and nuclear marker channel were converted to .TIFF format; 2) The DAPI channel image was utilized to generate individual ROIs for each nucleus in an image by background subtraction with a rolling ball radius of 50, auto-thresholding with the default algorithm, despeckling, nuclear segmentation using the watershed function, and finally generating ROIs via the Analyze Particles function with size 5 to Infinity and circularity 0.2 to 1.0; 3) The nuclear marker channel was then opened and background subtracted with rolling ball radius of 5. Mean intensities within the previously determined ROIs were measured with results output in a .CSV file; 4) .CSV files were qualitatively inspected with attention to the observed intensity distributions across GBOs of varying conditions; 5) Thresholds for assigning marker positivity were determined manually by measuring the mean intensity of nuclei with the minimal signal that would have been determined to be marker positive by traditional manual counting. The same threshold was applied across technical replicates under the same condition.

Cytoplasmic staining (cleaved caspase 3) quantification was performed as follows: 1) Zen 2 (.CZI) images for the DAPI and cytoplasmic marker channels were converted to .TIFF format; 2) the DAPI channel image was used to obtain the outline of the entire organoid by background subtraction with a rolling ball radius of 50, setting an extremely low manual threshold used across all images, and then generating a single large ROI corresponding to the entire organoid via the Analyze Particles function with size 5,000 to Infinity; 3) the cytoplasmic marker channel was then opened and background subtracted with rolling ball radius of 50. The mean gray value intensity of the single large ROI was measured with results output to a .CSV file.

Membrane staining (CD3, PD-L1, and EGFRvIII) quantification was performed as follows: CD3 staining was analyzed using the nuclear staining analysis pipeline as the CD3 immunoreactive cells were sparse, well circumscribed, and easily identifiable as individual cells; PD-L1 and EGFRvIII staining were analyzed using the cytoplasmic staining analysis pipeline as the signal was generally diffuse and individual cells were difficult to identify.

Statistical analyses were performed using the Student’s paired t-test with Bonferroni corrections for multiple hypothesis testing.

## ACKNOWLEDGMENTS

We thank all patients and their families who generously donated tissue, members of the Ming and Song laboratories and the Glioblastoma Translational Center of Excellence, Robert O. Heuckeroth, Zhaolan Zhou, Arjun Raj, Daniel Lim, and Kimberly M. Christian for their comments and suggestions, and J.G. Schnoll, B. Temsamrit, E. LaNoce, A. Angelucci, A. Garcia and G. Alepa for technical support and lab coordination. Some illustrations were created using Biorender. This work was partially supported by the Glioblastoma Translational Center of Excellence at the Abramson Cancer Center, Institute for Regenerative Medicine and Department of Neurosurgery at the University of Pennsylvania (to H.S., D.M.O., S.B. and M.N.), grants from the National Institutes of Health (R35NS116843 to H.S. and R35NS097370 to G-l.M.), Dr. Miriam and Sheldon G. Adelson Medical Research Foundation, and Pennsylvania Department of Health (to G-l.M.).

## AUTHOR CONTRIBUTIONS

D.Y.Z. led the study and performed the majority of the analyses. R.D.S. and D.Y.Z. optimized the co-culture model. D.R.C. and D.Y.Z. adapted and optimized the GoT method. R.T. generated CAR-T cells. X.W. and Y.S. performed immunostaining and imaging for the conditioned media experiments. Y.S. developed the image analysis pipeline and help with the writing. Q.C. and Y. Shi. contributed to the glioma cancer stem cell line analysis. B.H., R.D.S., F.J. and Z.Z. contributed to additional experiments and data collection, E.N. Jr., M.P.N., T.L., H.I.C., Z.A.B., S.B., R.D.S., and D.M.O. contributed to patient tissue collection. J.G. and J.W. contributed to T cell exhaustion analysis. D.Y.Z., G-l.M., and H.S. conceived the study and wrote the manuscript.

