## Supplementary Information for "Temporal multi-modal single-cell analyses reveal dynamic interactions of CAR-T cells with glioblastoma and targeting of antigen-negative neoplastic cells"

**Inventory**

Figures S1-7
Tables S1-7 legends
Supplementary Move 1 legend

SUPPLEMENTARY FIGURES

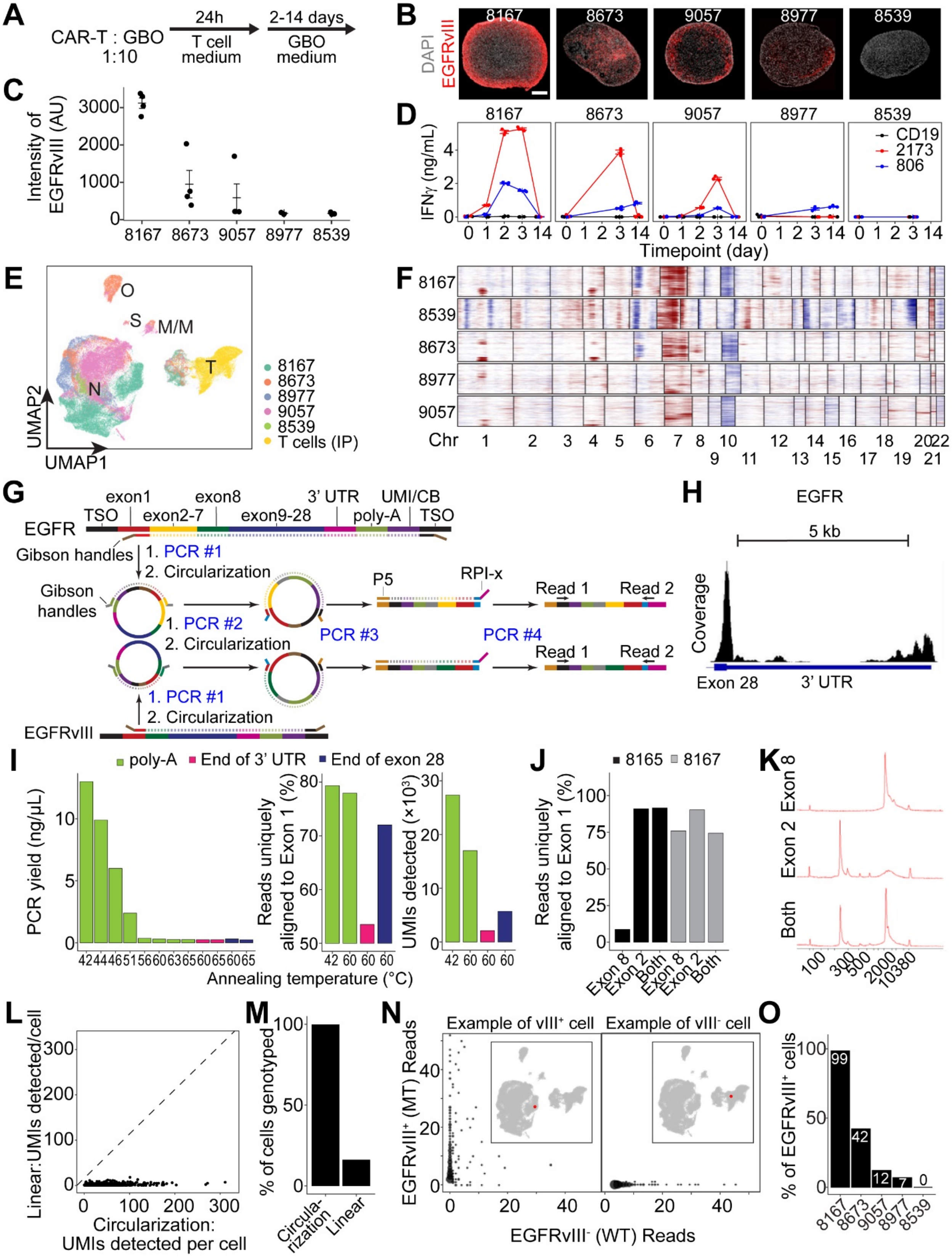


**Figure S1. EGFRvIII abundance in GBOs from different patients and optimization of single-cell EGFRvIII genotyping, related to Figure 1**

(A) A schematic diagram of the optimized CAR-T cell and GBO co-culture model. A ratio of 1 (CAR-T cells): 10 (GBO cells) in the cell number was used.
(B-C) Sample confocal images of EGFRvIII immunostaining in GBOs from 5 different patients used in the single-cell analyses (B) and quantification of EGFRvIII immunostaining intensity (C). Values represent mean $\pm$ SEM (n = 4).
(D) Quantification of IFNɣ concentration in the culture media by ELISA in the GBO and CAR-T cell co-cultures at different time points. Values represent mean $\pm$ SEM (n = 3).
(E) UMAP plot of the aggregated scRNA-seq data colored by different GBO and CAR-T cell co-cultures at all time points and the initial population (IP) of CAR-T cells.
(F) Heatmap of CNV inference analysis for neoplastic cells in GBOs from 5 different patients. Macrophages/microglia and T cell were used as the normal cell reference (not shown).
(G) A schematic diagram of the single-cell genotyping library preparation protocol describing intermediates generated from EGFRvIII mutant and wild-type transcripts. Targeted PCR #1 and #2 amplify regions flanking the exon 1-2 (wild-type) or 1-8 (EGFRvIII) junctions while preserving the cell barcode (CB) and UMI information. Circularization #1 brings the 5' end of the transcript into proximity to the CB and UMI that are located at 3' end of the cDNA molecule. Circularization #2 reverts the sequence inversion that occurs as a consequence of circularization #1. PCR #3 linearizes the targeted library, and PCR #4 adds Illumina sequencing adapters.
(H) Genome coverage plots of merged scRNA-seq reads at the 3' end of EGFR transcript in an EGFRvIII$^{+}$ GBO sample. Several distinct peaks were observed at the end of exon 28 and within the 5 kb long 3' UTR, indicating heterogeneity at the 3' end of the transcript, creating primer design challenges for the standard GoT method.
(I) Optimization of PCR #2 using different primers (oligo-dA, complementary to the end of the 3' UTR, and complementary to the end of exon 28) and different annealing temperatures. Shown are quantification of PCR #2 yield (left panel), percentages of reads uniquely aligned to exon 1 (middle panel) and number of unique UMIs detected (right panel).
(J) Optimization of PCR #3 using primers complementary to exon 2, exon 8, or both. Two samples were used, one without EGFRvIII and one with both EGFRvIII and wild-type EGFR. Shown is the quantification of percentages of reads uniquely aligned to exon 1.
(K) Bioanalyzer traces of libraries generated from samples without EGFRvIII transcripts using primers complementary to exon 2, exon 8, or both in PCR #3. The peak at ~2,000 bp corresponds to an insert spanning exons 1-8, and the peak at ~275 bp corresponds to an insert spanning only exons 1-2.
(L-M) Comparison of the circularization-based genotyping approach (G) with a direct linear enrichment approach, in which targeted PCR is performed just upstream to the exon 1 splice site to enrich for exon 1-2 (wild-type) and exon 1-8 (EGFRvIII) junctions. As shown in (L), far more UMIs per cell were identified by the circularization GoT based protocol. Shown in (M) is the quantification of the percentage of cells genotyped in the circularization-based approach vs. the direct linear enrichment approach.
(N) Examples of genotyping results for a prototypical EGFRvIII$^{+}$ cell and an EGFRvIII$^{-}$ cell. Overlapping data points are depicted by increased point sizes. Inset depicts a UMAP plot of scRNA-seq data with the cell highlighted.
(O) Quantification of the percentage of EGFRvIII$^{+}$ cells among all cells in each GBO based on genotype. Note that the genotyping result was very similar to those by immunostaining in (C).

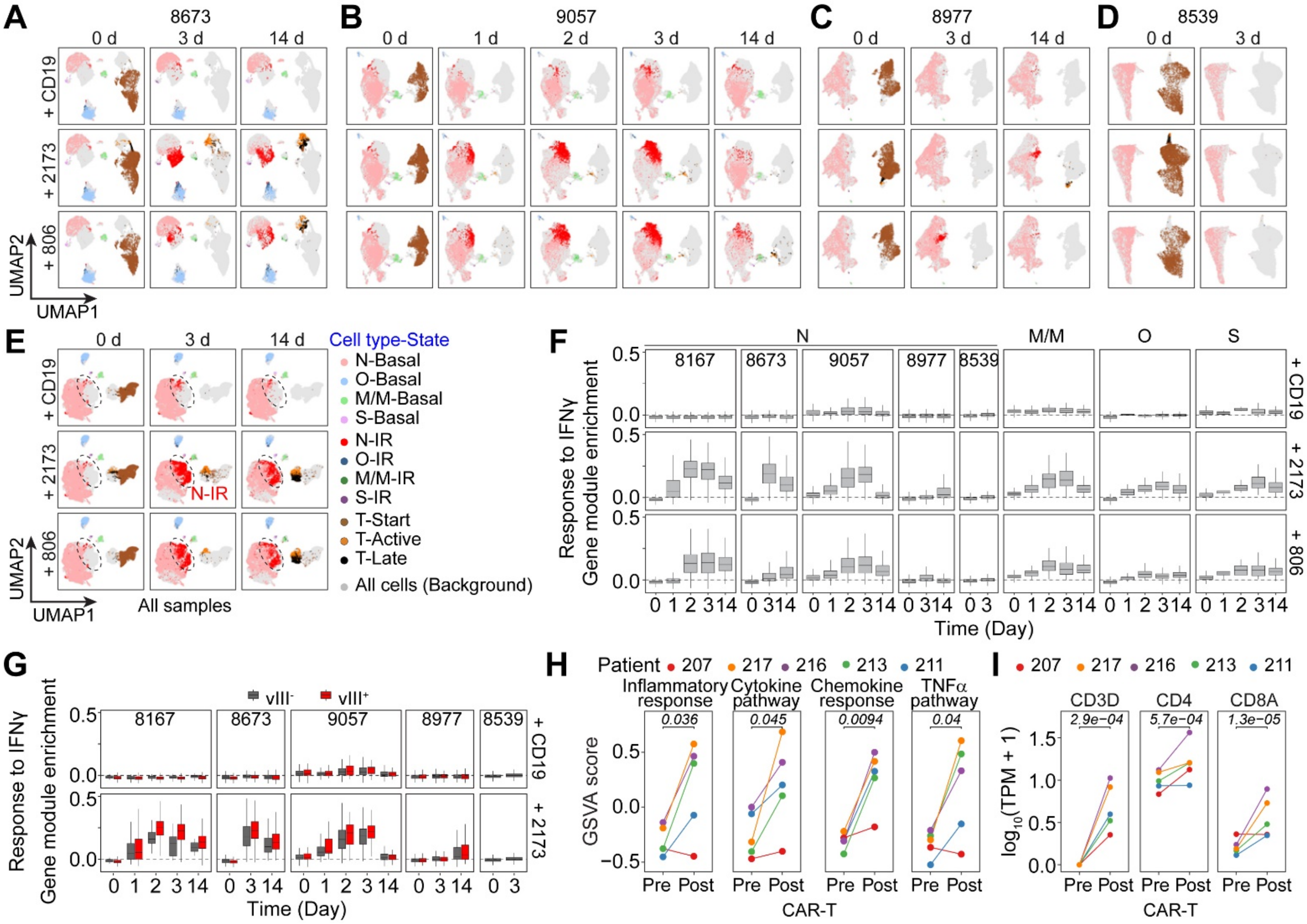


**Figure S2. Time-dependent shift of cell states in response to CAR-T cell treatment and analysis of IFNɣ response pathway enrichment, related to Figure 2**

(A-E) UMAP plots of scRNA-seq data of GBOs from individual patients co-cultured with CD19, 2173, and 806 CAR-T cells at different time points (A-D) and as well of aggregated data for GBOs from all different patients at all time points (E). Basal and IR states of different tumor cells, including neoplastic cells (N), oligodendrocytes (O), macrophages/microglia (M/M), and stromal cells (S), as well as start, active, and late states of T cells are indicated by different colors. The IR state of neoplastic cells is highlighted with a dashed line circle.

(F) Quantification of hallmark response to IFNɣ pathway enrichment of different cell types in different GBOs at different time points. Gene signature is taken from MSigDB[1].

(G) Quantification of IFNɣ response pathway enrichment of neoplastic cells of different GBOs separated by the EGFRvIII genotype. Note similar responses to 2173 CAR-T cells for EGFRvIII$^{+}$ and EGFRvIII$^{-}$ cells.

(H) GSVA scores of IR state enriched pathways in bulk RNA-seq data of pre- and post-CAR-T cell treatment patient samples from the 2173 trial. P-values were calculated using a paired student's t-test.

(I) Expression of T cell-associated genes in pre- and post-CAR-T cell treatment patient samples from the 2173 trial. Significance testing was performed using DESeq2 and Benjamini-Hochberg correction for multiple testing.

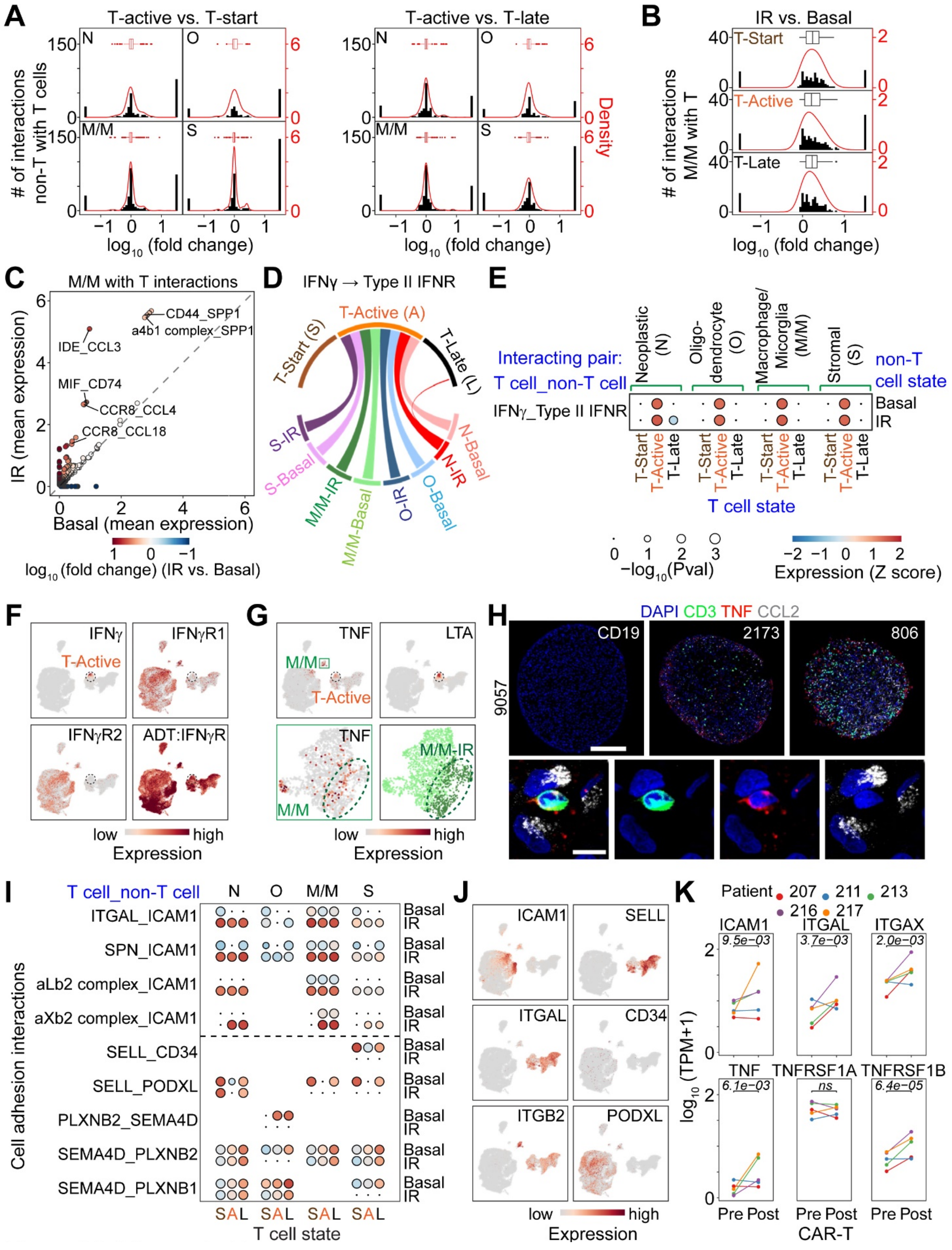


**Figure S3. Inferred cell-cell interactions between different T cell states and tumor cell states from ligand-receptor pair gene expression analysis, related to Figure 3**

(A) Histogram of the $\log_{10}$fold-change of the interaction strength between different tumor cell types with T cells of different states. Interactions that are not statistically significant in either IR or basal states are plotted at -1.5 and 1.5, respectively, for visualization purposes. Redline shows kernel density fit. Inset at the top shows a boxplot summarizing the distribution of differential interactions.
(B) Histogram of the $\log_{10}$fold-change of the interaction strength between IR vs. basal state macrophages/microglia with T cells of different states. Similar as in (A). Note that IR state macrophages/microglia have strengthened interactions with T cells from all states.
(C) Scatterplot of the interaction strength of T cells with macrophages/microglia between the basal and IR states. Point colors correspond to the $\log_{10}$fold-change of the strength of IR vs. basal states.
(D) Chord diagram of IFNɣ and its receptor Type II IFNR interaction between T cells and different tumor cells in different states. Width of chords correspond to the interaction strength.
(E) Bubble plot of IFNɣ and Type II IFNR interaction. Left hand y-axis indicates the specific ligand-receptor interaction. Upper x-axis indicates different tumor cell types. Right hand y-axis indicates basal vs. IR tumor cell state. Bottom x-axis indicates different T cell states. The bubble at the intersection of these four axes describes the specified interaction. Size of the bubble corresponds to the $-\log_{10}$P-value of the interaction as calculated by CellPhoneDB and the color indicates the interaction strength z-score across the same interaction between different pairs of cell types/states.
(F-G) UMAP plots of scRNA-seq data colored by mRNA or cell surface protein (ADT) expression as indicated. In (F), the T-active state is highlighted with a dashed line circle. Note that IFNɣ (F) as well as TNFα and LTA (G), is expressed only in active T cells, whereas IFNɣ receptor is expressed in all tumor cells at both the RNA and protein levels. Also shown in (G) are UMAP plots of subclustered macrophages/microglia highlighting the expression of TNF in the IR state (marked by a dashed line circle).
(H) Sample confocal images of CCL2 and TNF immunostaining of 9057 GBOs cultured with CD19, 2173, or 806 CAR-T cells for 3 days. Right-most images show a magnified view of $TNF^{+}CD3^{+}$ T-cells in proximity to $CCL2^{+}CD3^{-}$ tumor cells. Large scale bar: 250 μm, and small scale bar: 10 μm.
(I) Interaction bubble plot of cell adhesion-related interactions between T cells and different tumor subtypes. IR state enriched interactions are shown above the dashed horizontal line, and basal state enriched interactions are shown below.
(J) UMAP plots of scRNA-seq data colored by mRNA expression. Note that ICAM1 was upregulated in IR state neoplastic cells, whereas LFA1 (heterodimer of ITGAL and ITGB2) was expressed in all T cell states. The opposite pattern is seen with L-selectin and ligands CD34 and PODXL.
(K) Gene expression of ligand and receptor molecules in patient samples pre- and post- CAR-T cell therapy from the 2173 trial. Significance testing was performed using DESeq2 and Benjamini-Hochberg correction for multiple testing.

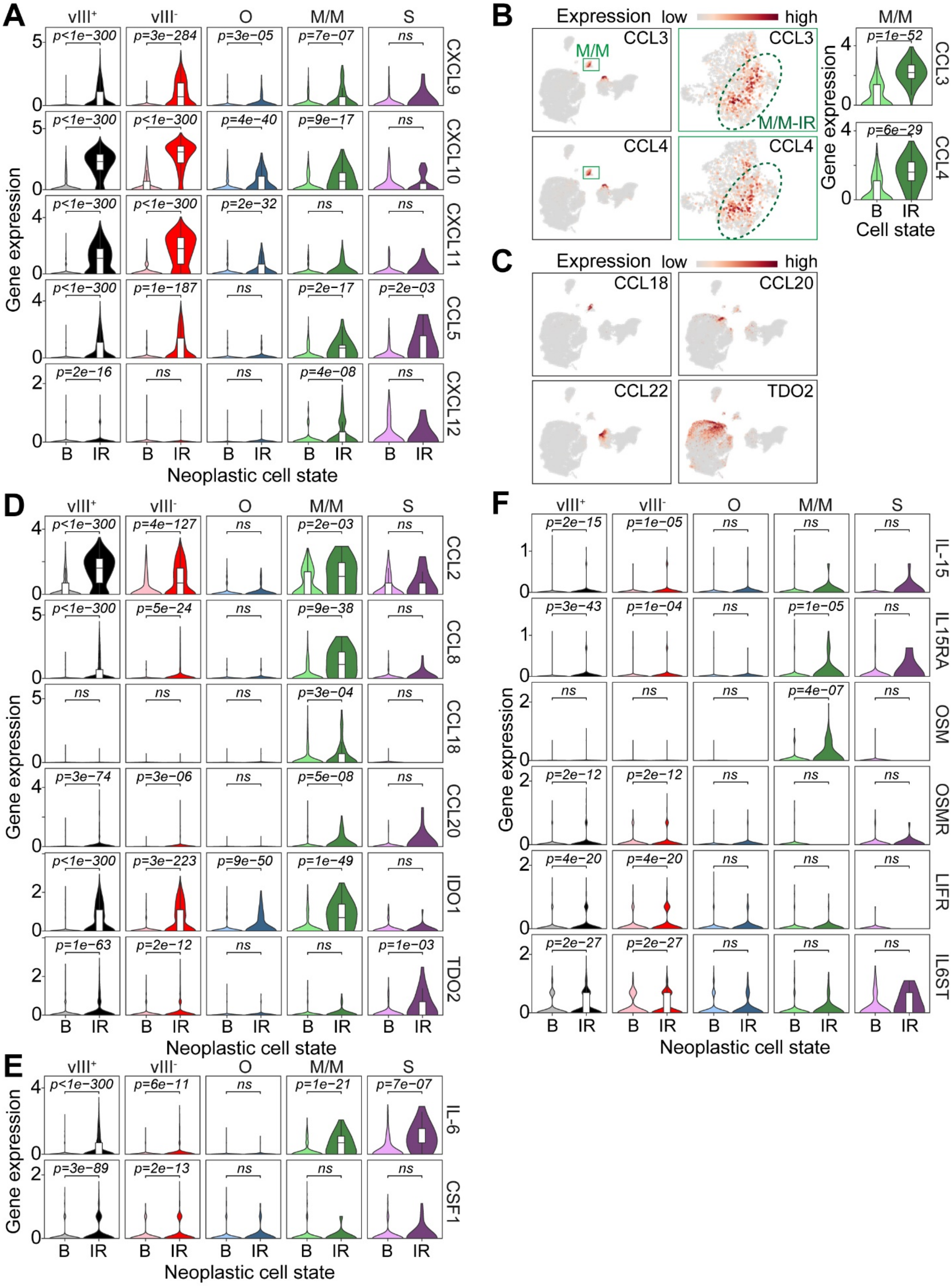


**Figure S4. CAR-T cell treatment induced changes in ligand-receptor interactions that can**

**recruit leukocytes, promote Treg differentiation, or promote M2 polarization of macrophages, related to Figure 4**

(A) Violin plots of expression of upregulated T cell recruitment chemokines in EGFRvIII$^+$ and EGFRvIII$^-$ neoplastic cells and in non-neoplastic cells. P values are calculated using the Wilcoxon rank sum test with Benjamini-Hochberg correction for multiple hypothesis testing.

(B) UMAP plots (left) and violin plots (right) of expression of T cell attracting chemokines, CCL3 and CCL4, upregulated in IR state macrophage/microglia. UMAP plots (middle) of scRNA-seq data just for the reclustered macrophages/microglia are also shown to highlight the expression in IR state (marked by a dashed line circle). See Figure S3G for the IR state of macrophages/microglia.

(C-D) UMAP (C) and violin plots (D) of expression of Treg recruiting chemokines (CCL2, CCL8, CCL18, CCL20, CCL22) as well as Treg differentiating metabolic enzymes (IDO1, TDO2) showing upregulation in tumor IR states.

(E-F) Violin plots of expression of upregulated M2 polarizing factors (IL-6 and CSF-1; E) and anti-tumor and pro-immune activity ligands and receptors (F) in EGFRvIII$^+$ and EGFRvIII$^-$ neoplastic cells and in non-neoplastic cells. Similar as in (A).

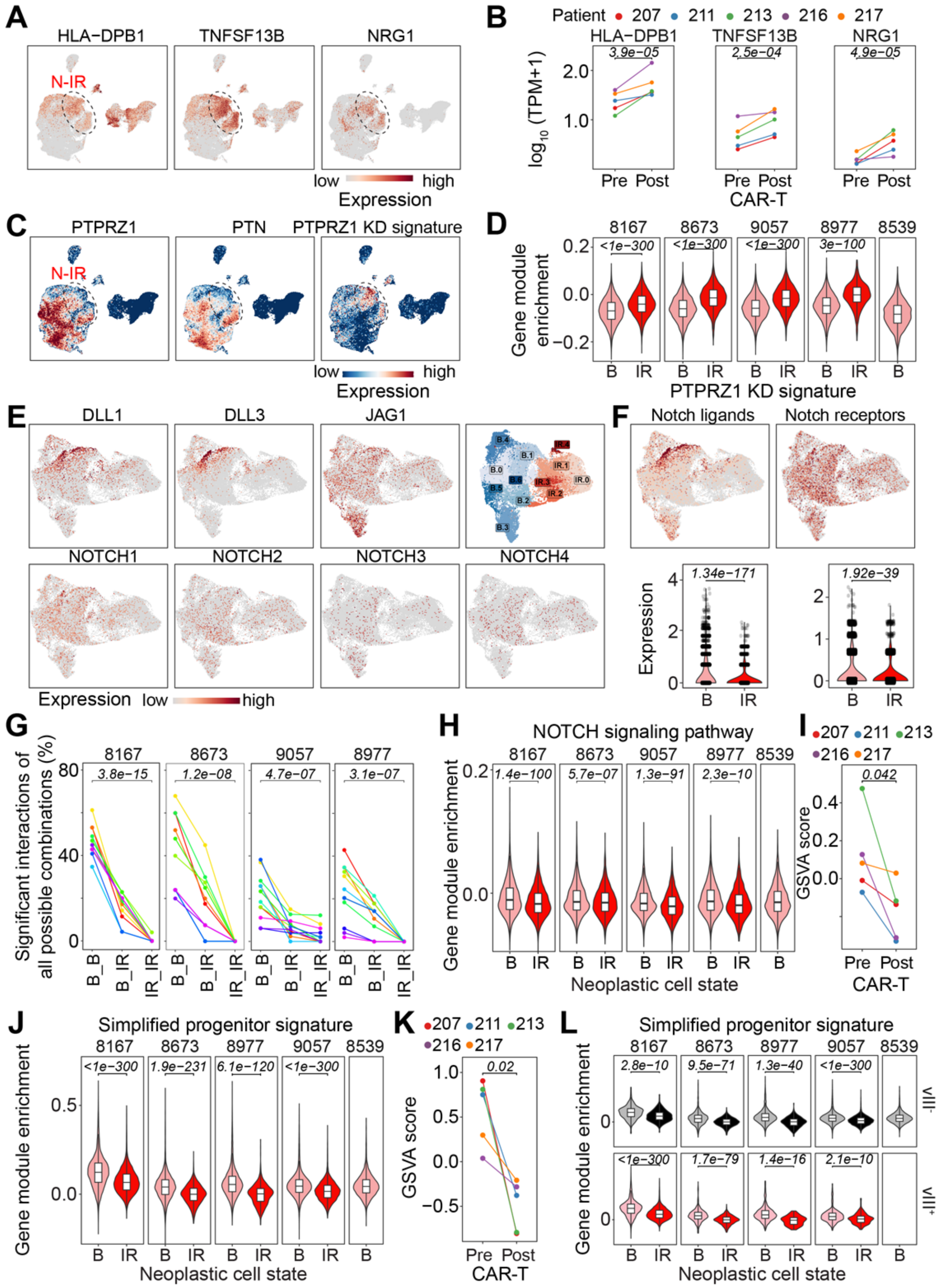


**Figure S5. CAR-T cell treatment induced changes in interactions within the neoplastic populations, related to Figure 5**

(A) UMAP plot of scRNA-seq data colored by expression of ligands and receptors corresponding to example upregulated interactions within neoplastic cell populations after the CAR-T cell treatment.
(B) Expression of factors as shown in (A) in pre- and post-CAR-T cell treatment patient samples from the 2173 trial. Significance testing was performed using DESeq2 and Benjamini-Hochberg correction for multiple testing.
(C) UMAP plot of scRNA-seq data colored by expression of PTPRZ1, PTN, and a PTPRZ1 knock-down signature from[2].
(D) Gene module enrichment for the PTPRZ1 knockdown signature in the basal vs. IR state of neoplastic cells. P values were calculated using a Wilcoxon rank sum test.
(E-F) UMAP and violin plots of expression of individual (E) and combined (F) Notch receptor and ligand molecules. The same UMAP plot of sub-clustered neoplastic cells as in Figure 5B is shown for reference.
(G) Quantification of Notch interactions within the basal state populations, between the basal and IR state populations, and within the IR state population of neoplastic cells as a percentage of all possible number of interactions given the subclustering. Statistical significance was calculated using one-way ANOVA.
(H) Gene module enrichment of Notch signaling pathway in the basal vs. IR state neoplastic cells.
(I) GSVA score of Notch signaling pathway in pre- and post-CAR-T cell treatment patient samples from the 2173 trial. P-values were calculated using a paired student’s t-test.
(J) Gene module enrichment for the simplified progenitor signature from[3] in the basal vs. IR state neoplastic cells.
(K) GSVA score of simplified progenitor signature from[3] in pre- and post-CAR-T cell treatment patient samples from the 2173 trial. P-values were calculated using a paired student’s t-test.
(L) Gene module enrichment for the simplified progenitor signature from[3] in the basal vs. IR state neoplastic cells separated by the EGFRvIII genotype. Only cells from 2173 CAR-T cell treatment were included in this analysis.

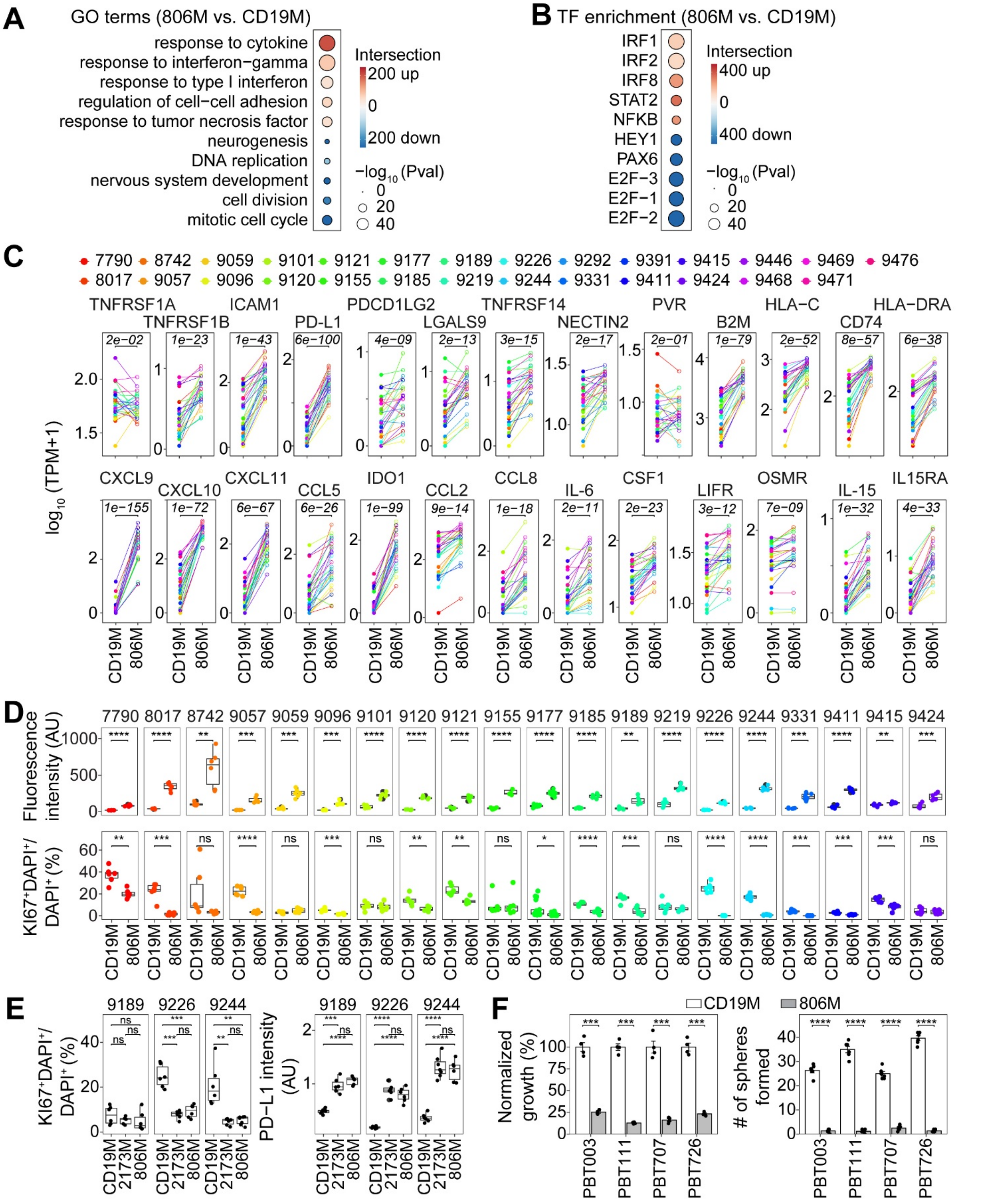


**Figure S6. Partial mimic of gene expression alterations observed in the IR state in GBO and CAR-T cell co-cultures by conditioned media, related to Figure 6**

(A-B) Bubble plots of up- and down-regulated GO terms (A) and transcription factor enrichment (B) in up- and down-regulated genes in 806M vs. CD19M treatment of GBOs derived from 27 patient

samples. Point size indicates -$\log_{10}$P-value and color indicates number of genes intersecting with the indicated pathways.
(C) Expression of genes listed in Figure 6E shown for GBOs derived from 27 individual patient samples. P-values were calculated using a paired student's t-test.
(D) Boxplots of quantification of PD-L1 immunostaining intensity and the percentage of $KI67^+$ cells shown for individual samples of GBOs derived from 20 different patient samples (n = 6; ns: $p \geq 0.05$; *p < 0.05; **p < 1e2; ***p < 1e3;****p < 1e4; paired student's t-test).
(E) Boxplots of quantification of PD-L1 immunostaining intensity and the percentage of $KI67^+$ cells shown for individual samples of GBOs derived from 3 different patient samples treated with CD19M, 2173M or 806M (n = 6; ns: $p \geq 0.05$; *p < 0.05; **p < 1e2; ***p < 1e3; ****p < 1e4; paired student's t-test).
(F) Barplots of cell growth and sphere formation in four different glioma stem cell lines exposed to CD19M or 806M. Values represent mean $\pm$ SEM (n = 3 for growth, n = 6 for sphere formation, ***p < 1e3; ****p < 1e4; student's t-test).

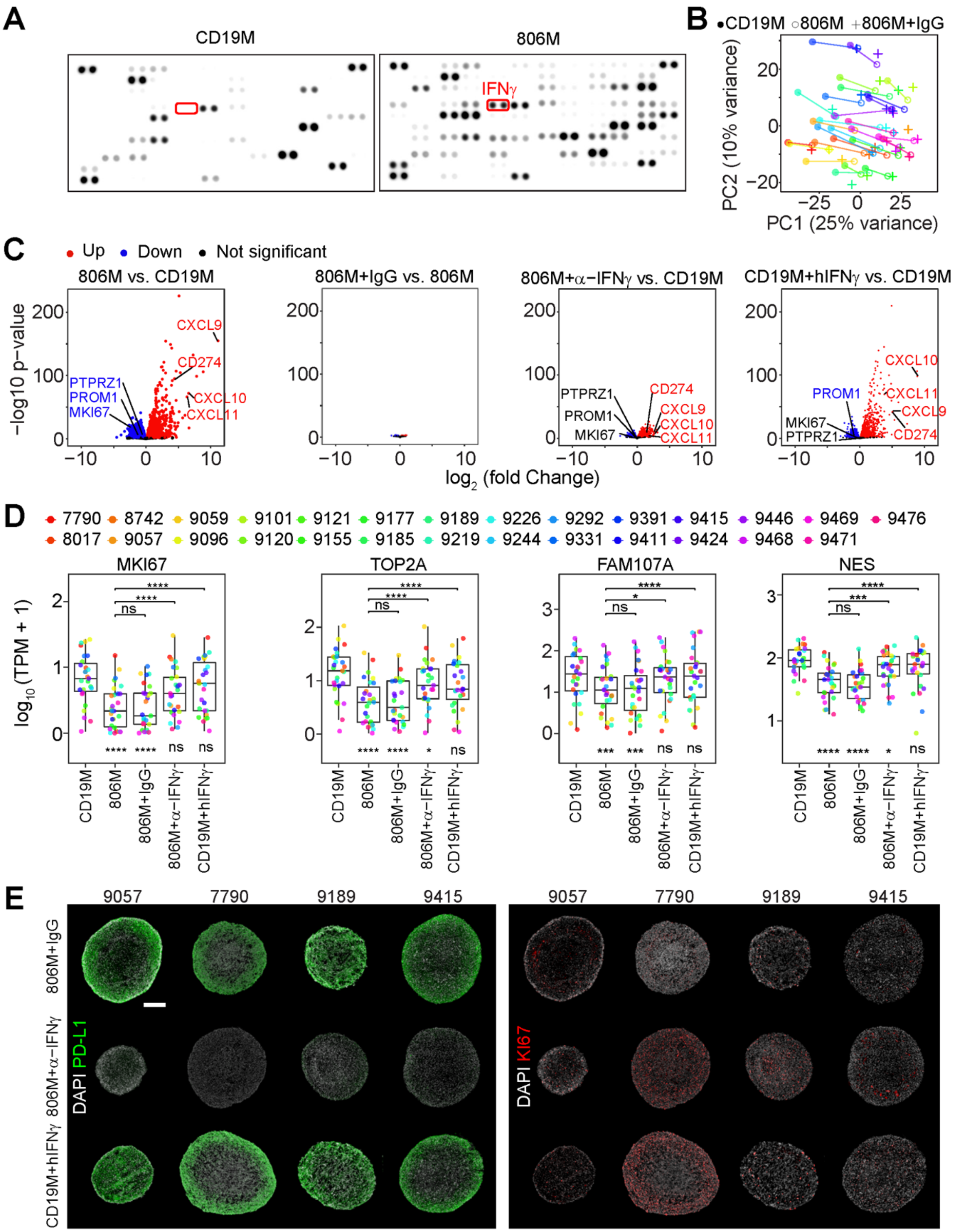


**Figure S7. Requirement, but not sufficiency of IFNɣ in mediating effects of conditioned media from GBO and CAR-T cell co-culture on tumor cells, related to Figure 7**

(A) Sample images of cytokine array membranes for CD19M and 806M with the IFNɣ spots highlighted.

(B) PCA plot of 806M with isotype control mouse IgG2A. CD19M and 806M points (same as in Figure 6A) are shown for reference.

(C) Volcano plots of 806M vs. CD19M (same as in Figure 6B, replotted as a reference), 806M with IgG vs. 806M, 806M with anti- IFNɣ vs. CD19M, and CD19M supplemented with IFNɣ vs. CD19M. Plots are shown with the same scales.
(D) Boxplots of quantification of expression of cell cycle and oRG markers with different conditioned media treatments. Asterisks below the boxplots and without brackets indicate statistical comparisons with CD19M. Asterisks with brackets correspond to statistical comparisons as indicated (ns: $p \geq 0.05$; *$p < 0.05$; **$p < 1e2$; ***$p < 1e3$;****$p < 1e4$; paired student's t-test).
(E) Sample confocal images of immunostaining for PD-L1 and KI67 in GBOs from 4 additional patients treated with 806M with IgG, 806M with anti-IFNɣ, and CD19M with IFNɣ. Scale bars: 200 $\mu$m.

### SUPPLEMENTARY TABLE LEGENDS (EXCEL FILES)

**Table S1. Patient sample information, related to Figures 1, 3, and 6**
List in (**A**) is summary of patient ID, age, sex, histological diagnosis, de novo or recurrent status, MGMT promoter methylation status, CPD (Center for Personalized Diagnostics at University of Pennsylvania) disease-associated mutation analysis panel, IDH status, and fusion transcript panel information for all patient samples. Single-cell analysis (**B**), 2173 trial (**C**), and conditioned media experiment samples (**D**) are listed individually.

**Table S2. Summary of information for multi-modal single-cell analyses, related to Figure 2**
List in (**A**) is summary of sample characteristics (patient ID, timepoint, and CAR-T cell treatment) and scRNA-seq features (# cells, average reads per cell, average genes detected per cell, and average UMIs detected per cell). List in (**B**) is a summary of Total-seq™ antibodies (Biolegend) used with their target protein, catalog #, and barcode sequence for CITE-seq analysis. List in (**C**) is a summary of oligos used in the EGFRvIII genotyping experiments.

**Table S3. List of differentially expressed genes in the tumor cell IR states, related to Figure 2**
Differentially expressed genes in IR states of neoplastic cells from different patients and aggregated non-neoplastic cells of all GBOs from scRNA-seq.

**Table S4. Summary of gene signatures used in the current study and their sources, related to Figure 2**

**Table S5. Summary of cell-cell interaction strengths and statistical significance, related to Figures 3, 5, and 7**
Listed are interaction strengths and statistical significance of T cells with (**A**) neoplastic cells, (**B**) oligodendrocytes, (**C**) macrophage/microglia, and (**D**) stromal cells, and interaction strengths and statistical significance of macrophage/microglia with (**E**) neoplastic cells, (**F**) oligodendrocytes, and (**G**) stromal cells.

**Table S6. List of differentially expressed genes in bulk RNA-seq from conditioned media experiments with log transformed fold-change and P-values listed, related to Figure 6**

**Table S7. Summary of cytokine array results, related to Figure 7**
Fold-change in the signal intensity between 806M and CD19M across both biological replicates.

### SUPPLEMENTARY MOVIE LEGEND

**Supplementary Video 1.** Live imaging of GBO and CAR-T cell co-culture. Time-lapse imaging of reaggregated 9057 GBOs (blue) with 806 CAR-T cells (red). Scale bar: 100 μm. Intervals between frames are 5 minutes. Movie also includes frames of a zoomed-in view with activated caspase 3/7 is marked in green, revealing CAR-T cell-induced cell death of 9057 GBO cells.
